# The Multifaceted Phenotype of Senescent HL-60/S4 Macrophages

**DOI:** 10.1101/2024.06.15.598082

**Authors:** Ada L. Olins, David Mark Welch, Dominik Saul, Igor Prudovsky, Donald E. Olins

## Abstract

Every cell has a multifaceted phenotype. Transcriptional analysis of functionally defined groups of genes can provide insight into this phenotypic complexity. In the present study, the mRNA transcriptome of phorbol ester (TPA) differentiated HL-60/S4 macrophage cells was scrutinized using Gene Set Enrichment Analysis (GSEA), which evaluates the strengths of various cellular phenotypes by examining the enrichment of functionally different gene sets. Employing GSEA, we obtained supporting evidence that HL-60/S4 macrophages are senescent, probably a consequence of enriched TGFβ and NOTCH signaling transcripts. There appears to be a reduction of transcripts for heterochromatin, nucleosome formation, and chromatin remodeling phenotypes. In addition, despite upregulated oxidative stress gene transcription, we observed a reduction of DNA damage and repair transcripts. GSEA indicated that transcripts for autophagy, extracellular matrix, and inflammation/inflammasomes are enriched. We also observed that the HL-60/S4 macrophage is enriched for apoptosis gene transcripts, which may promote necrotic death by pyroptosis. The long-term goal of this research direction is to see whether this complex multifaceted phenotypic pattern is shared with other types of macrophages and to determine what mechanisms might exist to coordinate these phenotypic facets within a single cell.

## Introduction

Cellular senescence is a state of irreversibly blocked replication (cell-cycling), while displaying senescence biomarkers, such as increased β-galactosidase, and generally exhibiting an absence of apoptotic death (Herranz and Gil, 2018; Hu et al., 2022; Huang et al., 2022; Rodier and Campisi, 2011; Soto-Gamez et al., 2019). Cellular senescence is distinguishable from “cellular quiescence”, where cell-cycle arrest is reversible (Marescal and Cheeseman, 2020), and from “terminal cell differentiation”, where cell-cycle arrest is accompanied by acquired tissue-specific functions (Li and Kirschner, 2014). Some terminally differentiated cells can become senescent, which is thought to prevent subsequent de-differentiation (Ring et al., 2022).

HL-60 cells, obtained from a human female with apparent Acute Promyelocytic Leukemia (APL), are capable of continuous growth in cell culture (Collins et al., 1977). In subsequent studies, HL-60 cells were shown to be Acute Myeloblastic Leukemia (AML), lacking the APL characteristic t(15:17) chromosome translocation. Also, they were demonstrated to be capable of *in vitro* cell differentiation (Collins, 1987). Retinoic acid (RA) induces the differentiation of HL-60 cells to granulocytes (neutrophils), while 12-O-Tetradecanoylphorbol 13-acetate (TPA or phorbol ester) induces their differentiation into macrophages. Subline HL-60/S4, the focus of the present study, was derived from HL-60 cells (Leung et al., 1992). *In vitro* differentiation of HL-60/S4 cells occurs about twice as rapidly, as in the parent HL-60 cell line.

In order to understand the HL-60/S4 differentiation phenotypes, we previously determined the polyA-mRNA transcriptomes of undifferentiated (0), granulocyte (RA treated) and macrophage (TPA treated) differentiated cells (Mark Welch et al., 2017). Initially, these data were mapped against the human genome (hg19) and we included all the splice variants for each gene (Mark Welch et al., 2017). In the present study, the same mRNA data are mapped against the human genome (hg38) and all splice variants for <u>each</u> gene are combined, yielding only “promoter-centered” transcripts for each gene (Table S1). In this study, these “promoter-centered” transcripts were employed to explore the differentiated macrophage cell state, compared to undifferentiated cells. Evidence is presented that TPA differentiated HL-60/S4 macrophages are senescent.

Cellular senescence is not a “single” cell state. The complexity of defining this cell state derives from multiple intrinsic and extrinsic factors, including cell-type and the nature of the cellular stresses. There are numerous types of senescence-inducing cellular stresses, for example: replicative failure; DNA damage; reactive oxygen species; oncogenic factors; NOTCH-inducing; inflammation, etc. The pathway to senescence for one kind of stress can differ, compared to the pathway of a different kind of stress. This senescence complexity is well known and described in numerous reviews (Fernández-Duran et al., 2022; Herranz and Gil, 2018; Hu et al., 2022; Huang et al., 2022; Rocha et al., 2022; Rodier and Campisi, 2011; Soto-Gamez et al., 2019).

Our observation that HL-60/S4 macrophages appear to be senescent is in agreement with studies which have shown that “primary” macrophages are also senescent (Behmoaras and Gil, 2021; Prattichizzo et al., 2016). The purpose of the present study is to characterize the HL-60/S4 macrophage multifaceted phenotype at day 4 of TPA treatment, compared to the undifferentiated cell state. This exploration encompasses many topics; e.g., nuclear and chromatin structure, cellular stress, autophagy, inflammatory response and mechanisms of cell attachment. Most of this exploration is based upon application of Gene Set Enrichment Analysis (GSEA) (Subramanian et al., 2005) on the available hg38 HL-60/S4 transcriptomes.

## Materials and Methods

### Cells Cultivation

HL-60/S4 cells can be purchased from ATCC (CRL-3306). They are cultivated in RPMI 1640 medium + 10% (unheated) Fetal calf serum + 1% Pen/Strep/Glut. We employ T-25 (6 ml) or T-75 (18 ml) flasks, lying horizontal to maximize surface area. The flasks are incubated humidified at 37° C with 5% CO_2_.

### Granulocytic Differentiation

Stock solution: 5 mM retinoic acid (RA) in 100% ethanol. We use Sigma-Aldrich R2625 (MW=300.4). Store the RA stock solution wrapped in aluminum foil at −20° C.

In a T-25 flask containing 5 ml of HL-60/S4 diluted with fresh medium to ∼2×10^5^ cells/ml, add 1µl of 5 mM RA. Wrap the flask with aluminum foil before placing in the incubator.

The cells will continue to replicate (more slowly) during the differentiation, gradually stopping. "Crawling" granulocytes begin to appear by day 3. After day 4, apoptosis becomes apparent. The cells never attach strongly to the flask bottom.

### Macrophage Differentiation

Stock solution: 80µM phorbol ester (TPA) in acetone. We use Sigma-Aldrich P1585 (MW=616.83). You can dissolve the TPA in absolute ethanol or DMSO. (1 mg dissolved in 20 ml). We use acetone, which means that you must pipette quickly. We do not use DMSO because it can induce granulocyte differentiation. Store the TPA stock wrapped in aluminum foil at −20° C.

The TPA treated cells become attached in one day (Figure 1). By day 3, fibroblast-shaped cells are apparent. By day 4, there are some dead cells in suspension. We aspirate the supernatant, then suspend the “live” cells using trypsin, followed by addition of fresh medium plus serum. We count the TPA cells with a hemacytometer. Typically, we find that the “live” cell concentration at day 4 is approximately equal to the concentration on day 0, indicating that the cells have stopped dividing early after addition of TPA.

**Figure 1.**
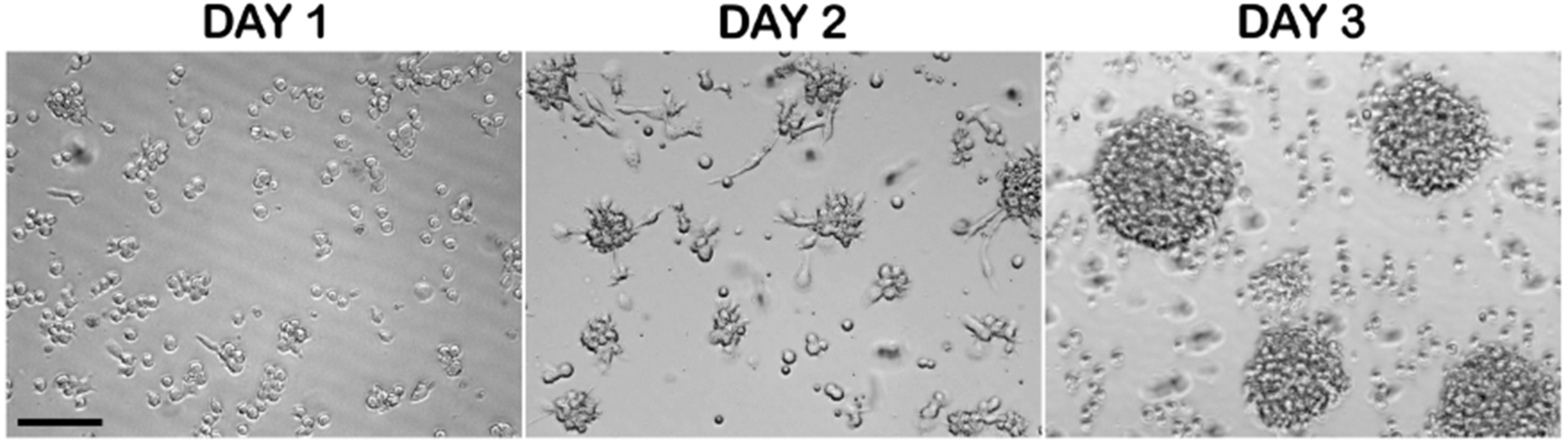
Phase microscopy of live HL-60/S4 cells incubating in 16 nM TPA. The cells are attached to the flask bottom by Day 1. They display formation of cell clumps with “migrating” fibroblast-type cells by Day 2, and often exhibit a “polka-dot” pattern of large cell clumps by Day 3. Day 4 looks identical to Day 3. “Day” refers to the days <u>after</u> addition of TPA. Magnification bar equals 100 µm. This figure is reproduced from (Mark Welch et al., 2017).

### Gene Set Enrichment Analysis (GSEA)

GSEA offers the opportunity to “match” an experimental <u>differential</u> gene expression list (e.g., TPA/0 or RA/0) against a curated GSEA gene set, compiled from functionally related genes (Mootha et al., 2003; Subramanian et al., 2005). The curated GSEA gene set is initially an unordered list. These curated genes become ordered (“ranked”) by comparison to their rank in the experimental differential gene expression list. Analysis of this ranked gene list documents the differential gene expression between the two cell phenotypes being compared (e.g., TPA versus 0). The Enrichment Score (ES) can be ES > 0.00 or < 0.00, reflecting the extent to which the curated gene set is overrepresented (+, upregulated) or underrepresented (-, downregulated) at the top or the bottom of the ranked list (e.g., TPA or 0). The nominal “p” value is an estimate of the statistical significance of an ES, where p = 0.00 is the maximal significance; p > 0.05 is regarded as less significant. The GSEA User Guide section on “Nominal P Value” describes that p-value accuracy can be increased by increasing permutations during the calculations (e.g., 100 to 1000 or more permutations). In most cases, we have employed 1000 permutations. On several occasions, when p = 0.06 or 0.07, we have employed increased permutations to explore whether the p values become closer to 0.05 (i.e., toward increased significance). Normalized Enrichment Score (NES) permits comparisons between different GSEA analyses, by accounting for variations in different gene set sizes. An important additional feature of the GSEA plot is identification of the “Leading Edge”, a subset (core) of the ranked gene set that contributes most to the ES. Analysis of the Leading Edge is a useful method for identification of the most influential genes in establishing a phenotype, which we have utilized quite frequently. The GSEA website (https://www.gsea-msigdb.org/gsea/index.jsp) explains how to conduct analyses and lists many functionally defined GSEA gene sets. Gene sets were chosen from the Molecular Signature Database (MSigDB), except for the SenMayo analysis (Saul et al., 2022).

As described earlier, the transcriptome data reported in 2017 (Mark Welch et al., 2017) was remapped to the hg38 assembly. The transcript levels are listed in Table S1. Data for the quadruplicate samples of each phenotype (i.e., 0, RA and TPA) were uploaded to the GSEA software (https://www.gsea-msigdb.org/gsea/index.jsp). GSEA produced “enrichment plots” and a list of the genes in the Leading Edge, which we present in this article to demonstrate the general results of these analyses. In some cases, we also present bar graphs showing the transcript levels (Log_2_FC) for relevant genes, changed during cell differentiation (i.e., RA/0 and TPA/0).

## Results

### HL-60/S4 Macrophage Cells exhibit Senescence

Figure 1 presents phase micrographs of HL-60/S4 cells incubated in 16 nM TPA for up to 3 days (day 4 looks identical to day 3). The transcriptomes of TPA and RA treated cells were obtained at day 4 for both cell states (see Table S1 for the hg38 “promoter-centered” transcriptomes).

A frequently employed, but imperfect, cytochemical “biomarker” for cellular senescence is the lysosomal β-galactosidase (β-gal) assay, which is generally performed at pH 6.0 on fixed cells. We employed a green fluorescent version of this assay (ThermoFisher Scientific “CellEvent Senescence Green”) to examine reactivity on formaldehyde-fixed HL-60/S4 0, RA and TPA cells (Figure 2). It is clear that TPA cells exhibit the strongest green fluorescence. This observation is in agreement with an earlier study (Yegorov et al., 1998), which argued that lysosomal β-gal “cannot be considered an exclusive marker of senescent cells, since it is expressed in other types of nonproliferating cells.” However, HL-60/S4 RA granulocytes have also ceased dividing by Day 4 (Olins et al., 1998), but show a weaker reaction than the TPA treated cells (Figure 2). The CellEvent Senescence Green data is “consistent” with TPA cells being senescent.

**Figure 2.**
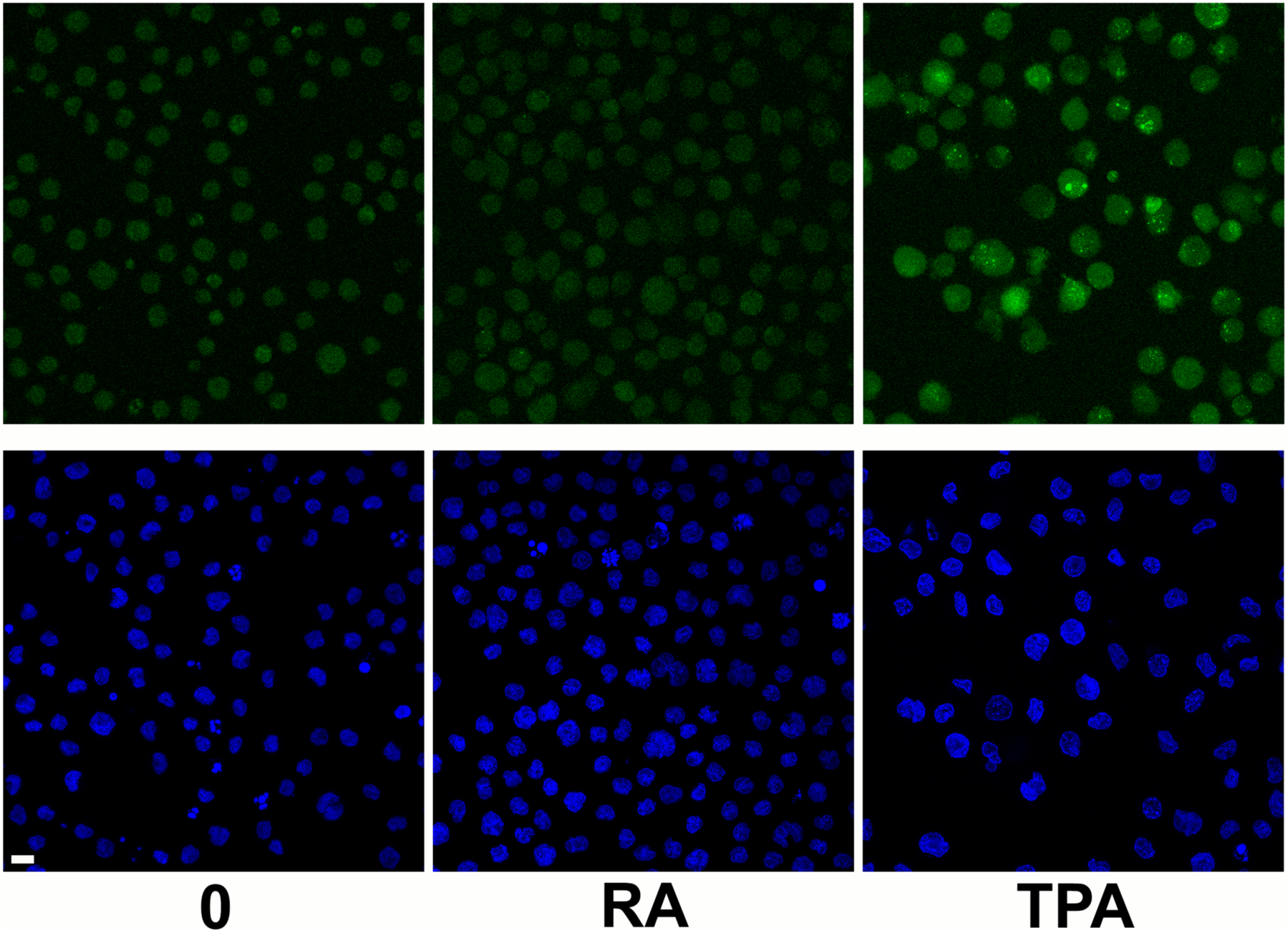
CellEvent Senescence Green Detection assay applied to formaldehyde-fixed HL-60/S4 0, RA and TPA cells. Top row: β-gal green fluorescence. Bottom row: DAPI blue staining of the identical “fields” of cells shown in the top row. Magnification bar equals 10 µm. Note that the TPA cells reveal stronger green fluorescence than the 0 and RA cells.

One of the principal characteristics of senescent cells is the cessation of cell division due to increased transcript levels of the cyclin-dependent kinase inhibitors (CDKN1A and CDKN2A). Cyclin-dependent kinase subunits (CDK2 and CDK4) are essential parts of the protein complex that controls G1 to S progression. Table 1 indicates that the transcript levels of CDKN1A and CDKN2A are especially elevated in TPA treated HL-60/S4 cells, compared to RA treated cells (based upon information in Table S1). In addition, Table 1 demonstrates that TPA treated cells exhibit a dramatic decrease in expression of the telomerase reverse transcriptase (TERT) gene. Although TERT expression was decreased in RA treated cells, this effect is more modest than after TPA treatment. Similarly, there was a very strong suppression of cyclin A2 (CCNA2, a key regulator of the cell cycle) gene expression in TPA treated cells, but less effect in RA-treated cells. These observations indicate that TPA treated cells are much more likely to be senescent, than are the RA treated cells.

**Table 1.** Comparison of the relative expression of cell cycle related genes in HL-60-S4 granulocytes (RA/0) and macrophages (TPA/0). Column Headings: Gene Name, common name; Code, gene code; Log_2_FC, log_2_ fold change of normalized gene expression levels; Inhibition, cell cycle enzymes inhibited.

| STABLE CESSATION OF THE CELL CYCLE |  |  |  |  |
| --- | --- | --- | --- | --- |
|  |  | Log<sub>2</sub>FC | Log<sub>2</sub>FC |  |
| Gene Name | Code | RA/0 | TPA/0 | Inhibition |
| p21 | CDKN1A | 3.78 | 10.68 | CDK2&4 |
| p16 | CDKN2A | -0.61 | 1.60 | CDK4 |
| Telomerase | TERT | -5.49 | -10.06 | -- |
| Cyclin A2 | CCNA2 | -0.58 | -3.46 | -- |

### Gene Set Enrichment Analysis (GSEA) Evidence of Senescence

The present GSEA employs a gene set (SenMayo) which was developed to assess the senescence-associated secretory phenotype (SASP) from various cells and tissues (Saul et al., 2022). Figure 3 clearly displays that the NES for TPA cells is greater than for RA cells, indicating that the TPA cells are more senescent, than the RA cells.

**Figure 3.**
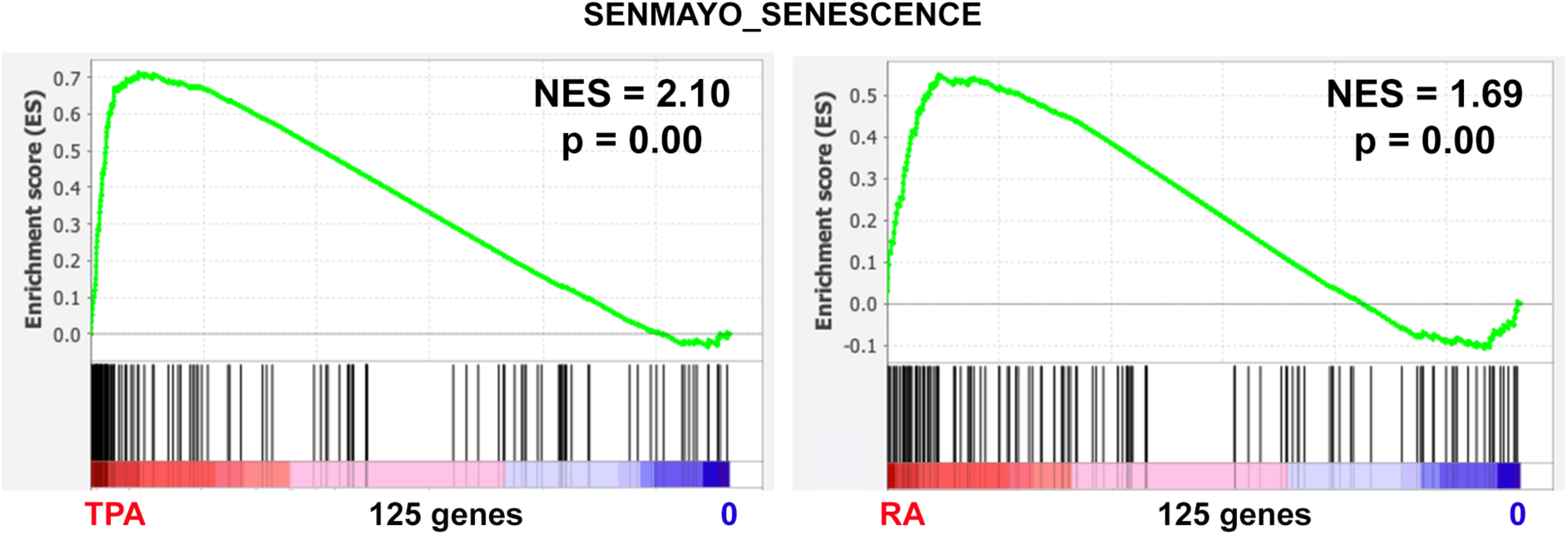
GSEA of genes expressed in HL-60/S4 cells, treated with TPA (left) or with RA (right). The 125 vertical black lines with the same height display the ranking and distribution of each member of the SenMayo gene set. For both the TPA/0 and RA/0 expression data, the SenMayo gene set is overrepresented at the cell differentiated end (i.e., TPA and RA, respectively), compared to the undifferentiated progenitor end (0). NES for TPA is greater than NES for RA, supporting that the TPA cells have a greater likelihood of being senescent, than the RA cells.

In summary, considering that TPA cells exhibit a “strong” β-gal fluorescence reaction, show high p21 and p16 expression and reveal more significantly enriched SenMayo senescence-related gene expression than the undifferentiated progenitor (0) cells, we conclude that TPA treated HL-60/S4 cells are most likely senescent. The remainder of this study will concentrate upon exploration of the phenotypic properties of these senescent TPA macrophages, primarily (but not exclusively) employing GSEA. Since RA treated cells are differentiated along an alternative pathway (i.e., granulocyte), with less indication of senescence, their phenotype will be less frequently discussed in this paper. However, their hg38 transcriptome (RA/0) can be found with the (TPA/0) transcriptome on separate sheets of Table S1.

### Changes in Nuclear Shape, the Nuclear Envelope and Cytoskeletal protein transcripts in Senescent HL-60/S4 Macrophage Cells

A number of excellent articles concerned with chromatin changes during senescence have been published (Olan et al., 2020; Parry et al., 2018; Parry and Narita, 2016; Rocha et al., 2022; Sikder et al., 2022; Sławińska and Krupa, 2021; Swanson et al., 2015). These articles document both the similarities and differences observed comparing cellular senescence in different types of cells. With this perspective, we briefly describe the varying microscopic phenotypes of the HL-60/S4 cell system: Undifferentiated (0) cells and granulocytes (RA) grow in suspension. Granulocytes are weakly associated with the substrate, but capable of “crawling”. As described earlier (Figure 1), TPA induced macrophages bind to the substrate and to each-other often forming cell clumps. The macrophage nuclei look very “distorted”, with little resemblance to the undifferentiated progenitor cell nuclei (Figure 4).

**Figure 4.**
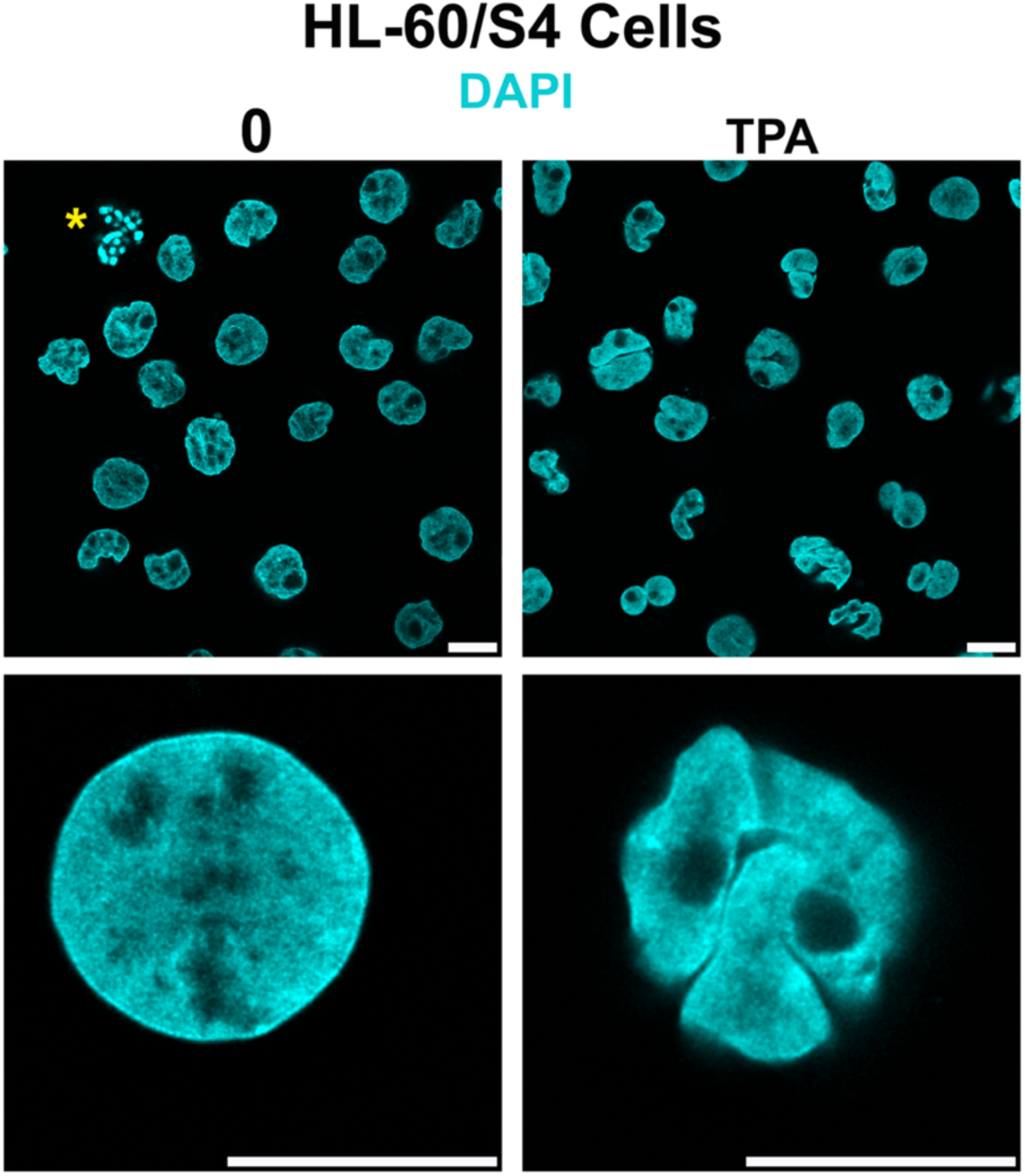
DAPI stained undifferentiated (0) and macrophage (TPA) HL60/S4 cells. Magnification bar equals 10 μm. The yellow asterisk highlights a near-by mitotic cell.

In HL-60/S4 cells, phenotypes are reflected in differences of the hg38 transcriptomes for protein components of the nuclear envelope (Figure 5) and the cytoskeleton (Figure 6), documenting earlier studies (Mark Welch et al., 2017; Olins et al., 2009; Olins and Olins, 2005; Rowat et al., 2013). The downregulation of LMNA (Lamin A) in RA-induced granulocytes appears to augment the deformability of the nuclear envelope (NE); upregulation of LMNA in HL-60/S4 granulocytes decreases the deformability (Rowat et al., 2013). The increase in SUN2 and SYNE2 (Nesprin-2) in macrophage, in combination with ACTB (β-Actin) likely enhances the LINC (Linker of Nucleoskeleton and Cytoskeleton) complex formation, strengthening the connections between the NE and the cell membrane, via the cytoskeleton (Bouzid et al., 2019). PLEC (Plectin) can also interconnect actin with intermediate filaments (e.g., VIM, Vimentin), further making the macrophage nucleus more stable in its positioning within the cell. In addition, SPTAN1 (Spectrin) is upregulated in TPA cells. This protein forms a scaffold stabilizing the cell membrane and can also cross-link actin (Machnicka et al., 2019).

**Figure 5.**
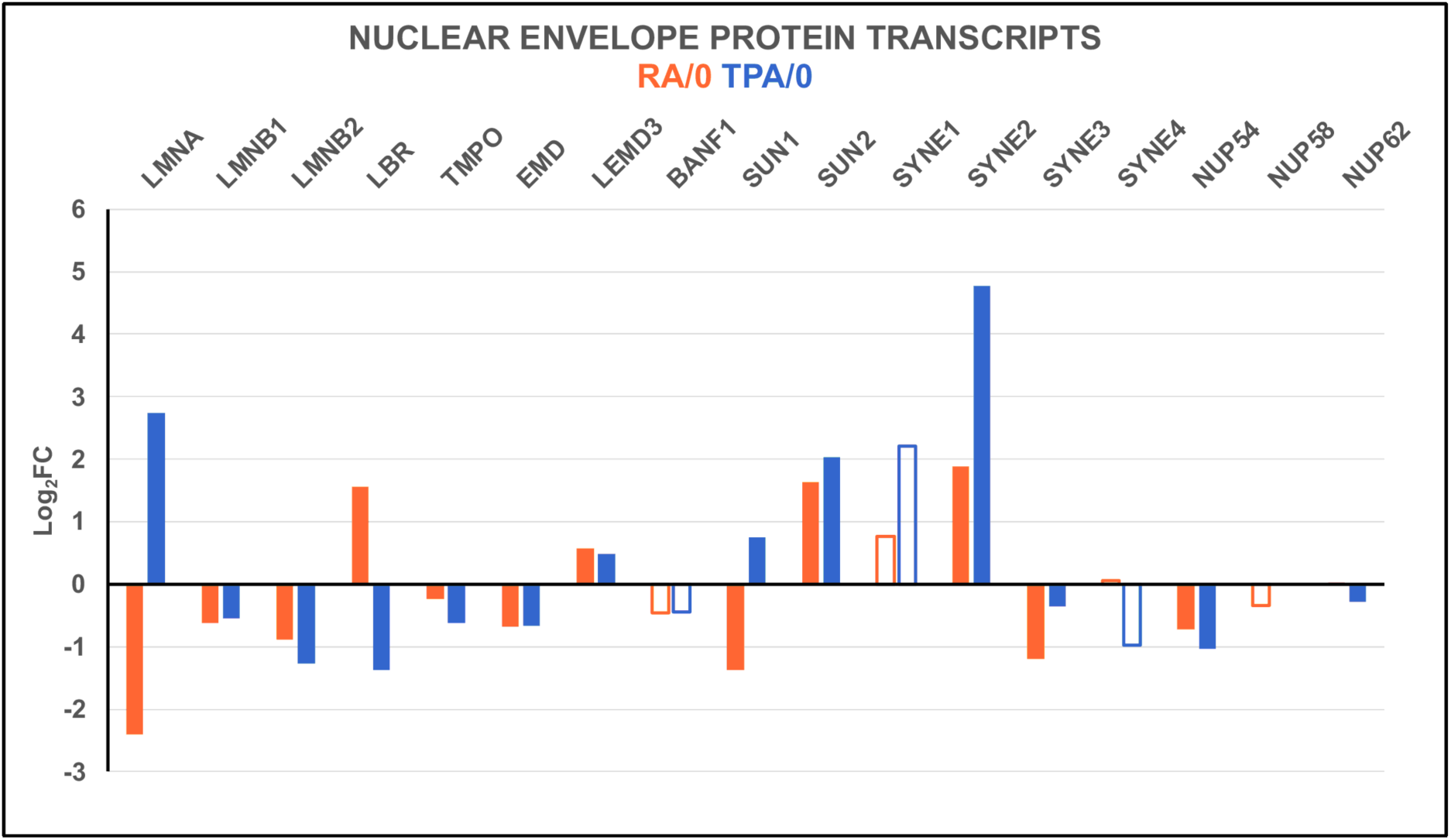
Nuclear Envelope protein Log_2_FC transcript levels. Note that LMNA is reduced in RA cells, making the granulocyte nucleus more deformable; but LMNA is elevated in TPA cells. LBR exhibits reciprocal changes, compared to LMNA. Also note the increases in SUN2 and SYNE2 (Nesprin-2) in TPA cells. The open bars indicate lower statistical significance (PPDE < 0.95); solid bars indicate a PPDE > 0.95.

**Figure 6.**
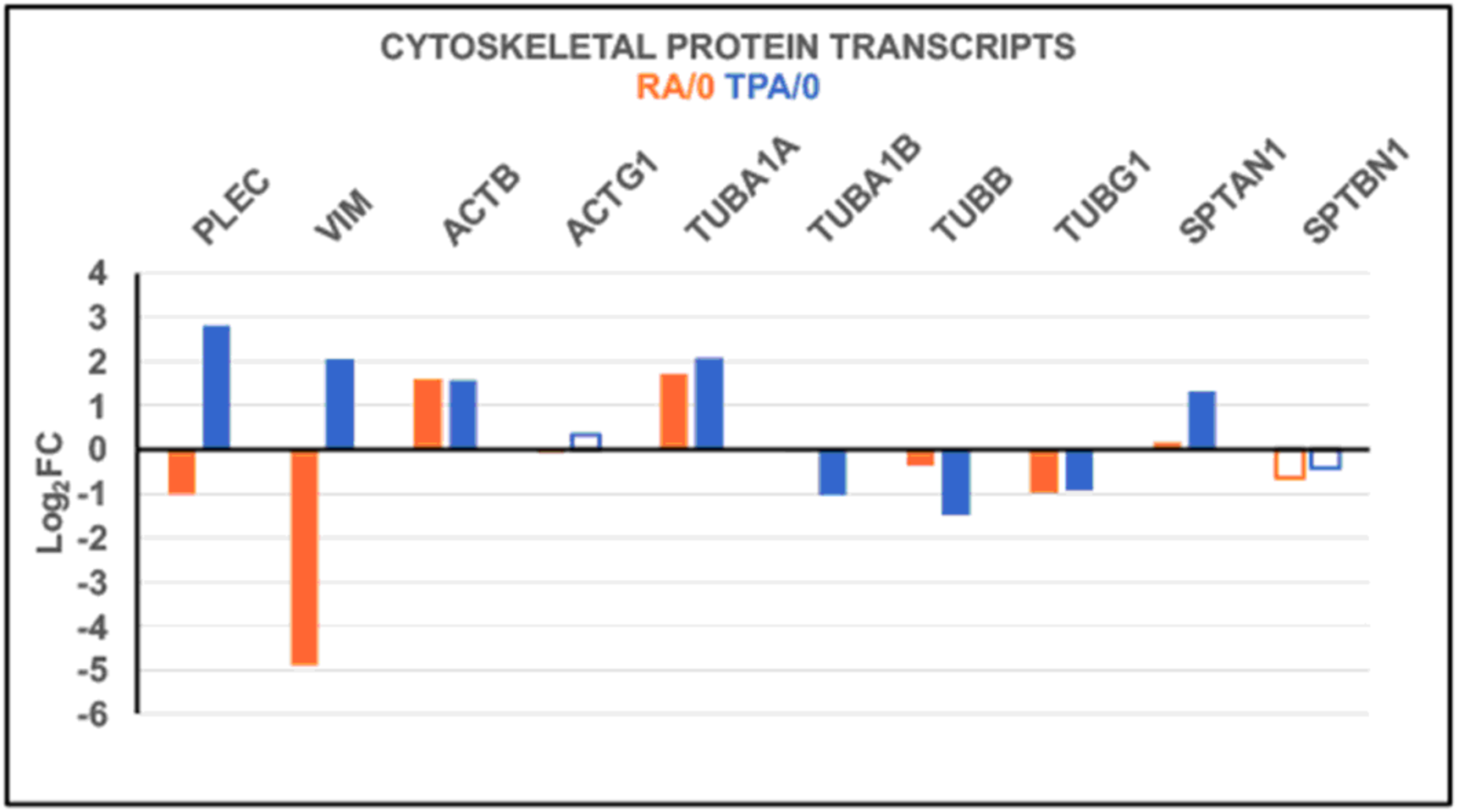
Cytoskeletal protein Log_2_FC transcript levels. Note that Plectin (PLEC) and Vimentin (VIM) are both downregulated in granulocytes and upregulated in macrophages. Note also that Alpha Spectrin-1 (SPTAN1) is elevated in macrophages, but not in granulocytes. This protein forms a scaffold stabilizing the cell membrane and can also cross-link actin. The open bars indicate lower statistical significance (PPDE < 0.95); solid bars indicate a PPDE > 0.95.

It should also be pointed out that a deficiency of LMNB1 (Lamin B1) is regarded as a biomarker of cellular senescence and considered causative for many changes in chromatin structure (Freund et al., 2012; Radspieler et al., 2019; Shah et al., 2013; Shimi et al., 2011). However, Figure 5 illustrates that LMNB1 is slightly less in RA cells, than in TPA cells. Perhaps LMNB1 transcript levels are not as meaningful to senescence causation in HL-60/S4. We propose that the combined decrease in LBR transcripts, and those of LMNB1 and LMNB2, in TPA cells may predispose HL-60/S4 cells for senescence, a suggestion consistent with prior publications (Castro-Obregón, 2020; En et al., 2020).

### Heterochromatin Changes in Senescent TPA treated HL-60/S4 Macrophage Cells

Numerous studies have presented evidence that senescent cells exhibit a reduction in global heterochromatin (Parry and Narita, 2016; Rocha et al., 2022; Sławińska and Krupa, 2021; Swanson et al., 2015; Zhang et al., 2021). Our goal was to determine whether this generalization accurately describes the macrophage chromatin of TPA treated HL-60/S4 cells. For this purpose, we chose to use Gene Set Enrichment Analysis. Four different heterochromatin GSEA gene sets were matched against the same TPA/0 expression data (Figure 7a). The four plots of the different GSEA analyses resemble each other, yielding comparable NES and nominal p values scores, despite employing different gene sets (i.e., differences in gene numbers and selection of genes). The negative NES values from all four gene sets indicates that the gene transcript levels responsible for establishing global heterochromatin are underrepresented (reduced) in the TPA differentiated macrophage. To identify these putative “heterochromatin” genes, we created a list and graph of the most prevalent gene names, comparing the Leading Edge genes from the four different GSEA gene sets (Figure 7b). Clearly, experiments are required to test whether these candidate genes either singly or in combination can facilitate heterochromatin formation or can diminish heterochromatin when these transcript levels are reduced.

**Figure 7.**
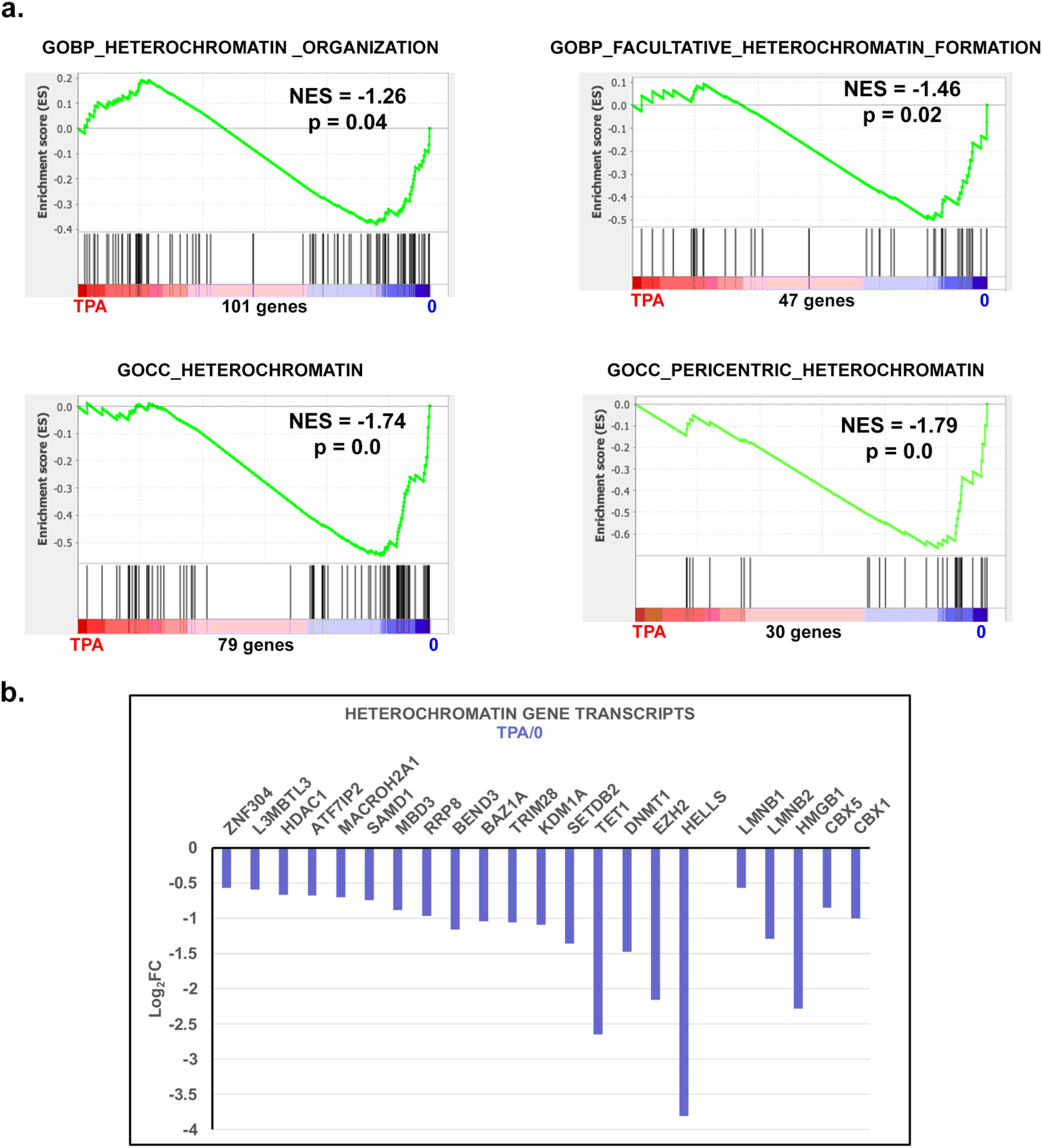
Heterochromatin Changes. a) GSEA plots with four different heterochromatin gene sets. The number of genes in each gene set are shown at the bottom of each plot. From the shape of these plots and the increased density of ranked genes at the undifferentiated (0) phenotype location, it is clear that most of the genes in each gene set are underrepresented in the macrophage phenotype region. We conclude that heterochromatin is reduced in TPA differentiated HL-60/S4 cells. b) Candidate heterochromatin forming genes. The genes presented (on the left side) of the Log_2_FC graph were “distilled” from the Leading Edge of each of the four GSEA heterochromatin analyses shown in 7a. For a gene to be presented, it had to be listed in two-or-more Leading Edges. Also shown (on the right side) are several genes that appear in only one Leading Edge, but are frequent candidates for heterochromatin formation (i.e., LMNB1, LMNB2, HMGB1, CBX5 and CBX1). All of the listed genes exhibited a PPDE > 0.95 and a Log_2_FC < 0.0

### Nucleosome Formation: Reduction of Transcripts in HL-60/S4 Macrophages

The decreases of heterochromatin within nuclei of senescent HL-60/S4 macrophages possibly reflects a changing composition of nucleosomes and nucleosome binding proteins. GSEA provides several gene sets focused at the nucleosome level. These gene sets include genes encoding histones (mostly, Replication Independent, RI), histone chaperones and other proteins involved with chromatin assembly. Figure 8a presents three different GSEA plots. It is clear from the negative NES values, that the bulk of genes in each gene set are not enriched in the TPA differentiated cells. Figure 8b presents Log2FC bar graphs of the GSEA Leading Edges from these plots. The genes presented represent a selection from the three Leading Edges, emphasizing those presented in <u>both</u> the GOBP and GOMF plots. Examples of significantly downregulated gene transcripts, including their GeneCards (genecards.org) functions are: CHAF1A and 1B, chromatin assembly and heterochromatin maintenance; H2AB1 and B3, RI histones; HMGA1 and B2, nucleosome phasing; MACROH2A1, RI histone, represses transcription and inhibits histone acetylation; NAP1L1 and L4, histone chaperones; H2AX, RI histone, DNA repair; H2AZ1, RI histone, heterochromatin formation. Note also, the apparently decreased levels of histone H1 transcripts.

**Figure 8.**
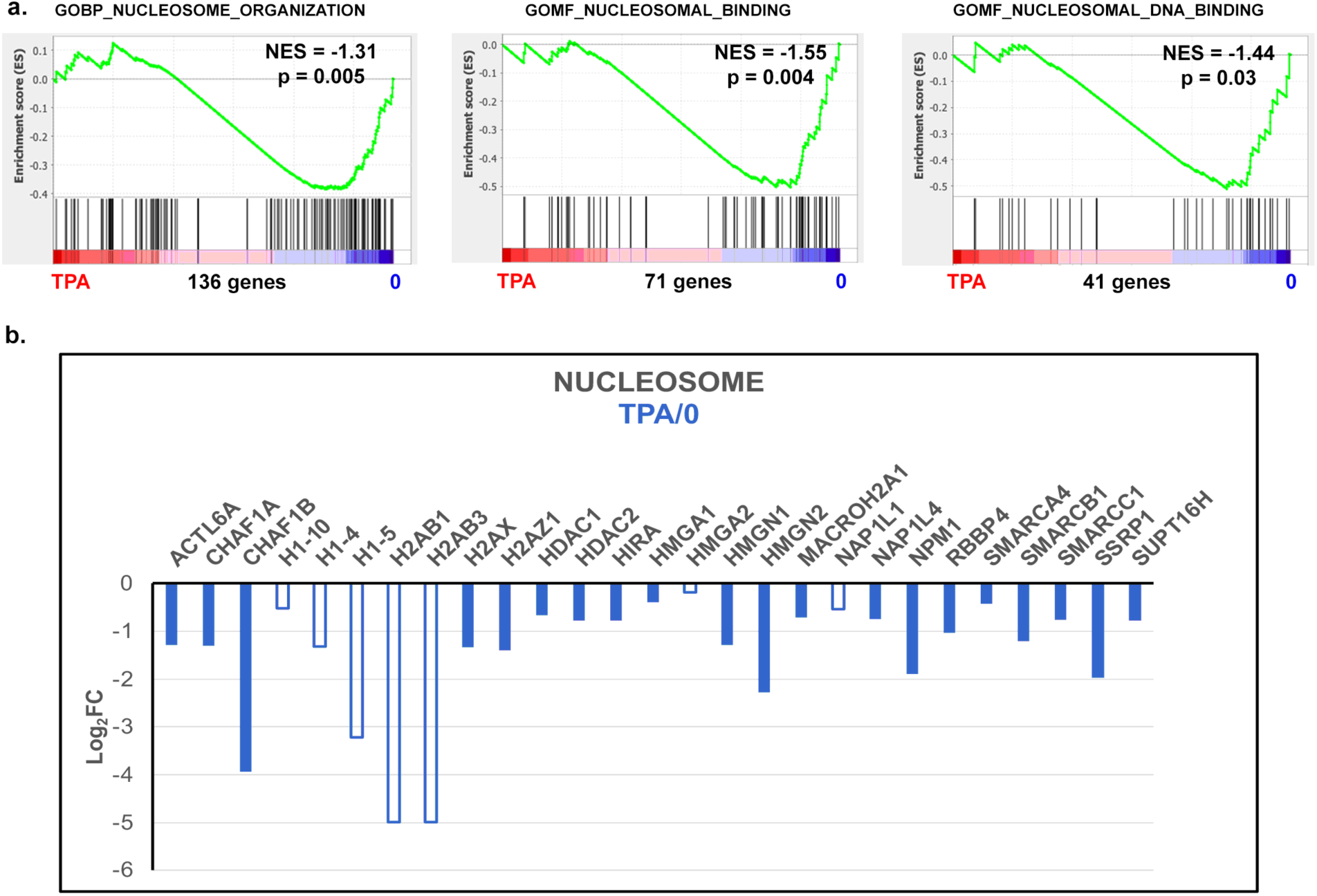
Nucleosome Formation. a) GSEA plots with three different nucleosome organization and binding gene sets. The number of genes in each gene set are shown at the bottom of each plot. From the shape of these plots and the increased density of ranked genes at the undifferentiated (0) phenotype location, it is clear that most of the genes in each gene set are underrepresented in the macrophage phenotype region. b) Genes presented in the Log_2_FC graph were selected from the Leading Edges of each of the three GSEA Nucleosome analyses, shown in 8a. The open bars indicate lower statistical significance (PPDE < 0.95); solid bars indicate a PPDE > 0.95.

### Chromatin Remodeling: Reduction of Transcripts in HL-60/S4 Macrophages

Cycling cells, such as undifferentiated HL-60/S4 0, have dynamic changes in gene expression during each cycle that involves machinery for moving nucleosomes to expose (or conceal) promoters, coding regions, etc. This provokes the question: Do senescent HL-60/S4 TPA treated macrophages also have the necessary chromatin remodeling machinery? GSEA has two gene sets on this subject: “Chromatin Remodeling” and “ATP Dependent Chromatin Remodeler Activity”. Figure 9a displays gene enrichment plots for these two gene sets, including NES and nominal p values. The results clearly demonstrate that for TPA compared to 0, the remodeling gene sets are underrepresented at the TPA end of the ranked gene list. In other words, TPA induced macrophages are deficient in the ATP-dependent and independent machinery for evicting or repositioning nucleosomes. Among the ATP-dependent remodeling activities, the SNF/SWI cluster is well studied (Centore et al., 2020). Figure 9b presents a Log_2_FC transcript summary of TPA/0, with regard to SNF/SWI protein subunit transcript levels, demonstrating minor differences in SMARCB1 and SMARCC1. It is of interest that SMARCB1 binding to the nucleosome acidic patch is necessary for proper remodeling (Centore et al., 2020). There are major downregulations of BCL7A and BCL11A, which are transcription factors for the highly studied BAF SNF/SWI remodeler complex. In summary, GSEA argues that TPA differentiated, senescent HL-60/S4 macrophages possess a significant reduction, compared to undifferentiated cells, in the transcripts for ATP-dependent or independent chromatin remodeling.

**Figure 9.**
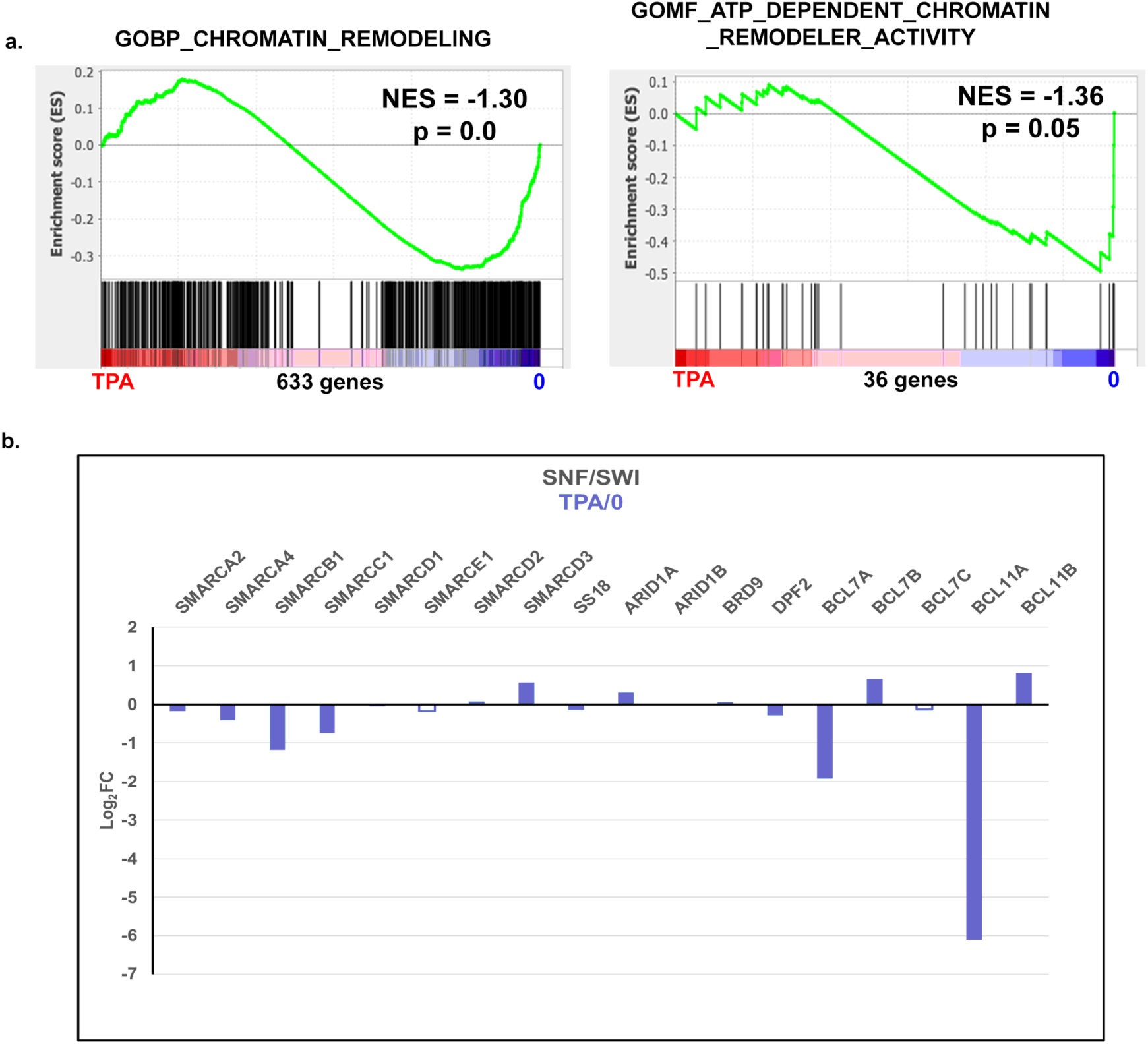
Chromatin Remodeling. a) GSEA plots of “Chromatin Remodeling” and “ATP-Dependent Chromatin Remodeling Activity”. b) Log_2_FC of SNF/SWI subunit protein transcripts in TPA/0 conditions. The open bars indicate lower statistical significance (PPDE < 0.95); solid bars indicate a PPDE > 0.95.

### mRNA Transcription: Upregulation in HL-60/S4 Macrophages

The reduction of heterochromatin in senescent HL-60/S4 macrophages (described earlier) implies that these nuclei have increased “open” chromatin, conducive for transcription. Indeed, GSEA has a single gene set on this subject: “GOBP_MRNA_TRANSCRIPTION”, which clearly shows that transcription augmenting factors are significantly enriched in the TPA-induced macrophage phenotype compared to the undifferentiated phenotype (Figure 10). However, increased nascent transcription is only part of the story. mRNA “processing”, including: 5’ capping, splicing and addition of a 3’ polyA tail must be completed before the “mature” mRNA can be exported from the nucleus into the cytoplasm (Hocine et al., 2010). It appears that some of the mRNA in the senescent HL-60/S4 macrophages are not adequately processed and unlikely to be translated. A GSEA plot with the gene set title “GOBP_MRNA_PROCESSING” illustrates the reduction of transcripts involved in mRNA processing. This leads to a reduction of transcripts being exported to the cytoplasm; see the GSEA plot with the gene set title “GOBP_MRNA_EXPORT_FROM_NUCLEUS”. Evidently, the SASP (Senescence Associated Secretory Phenotype) transcripts can circumvent this barrier into the cytoplasm.

**Figure 10.**
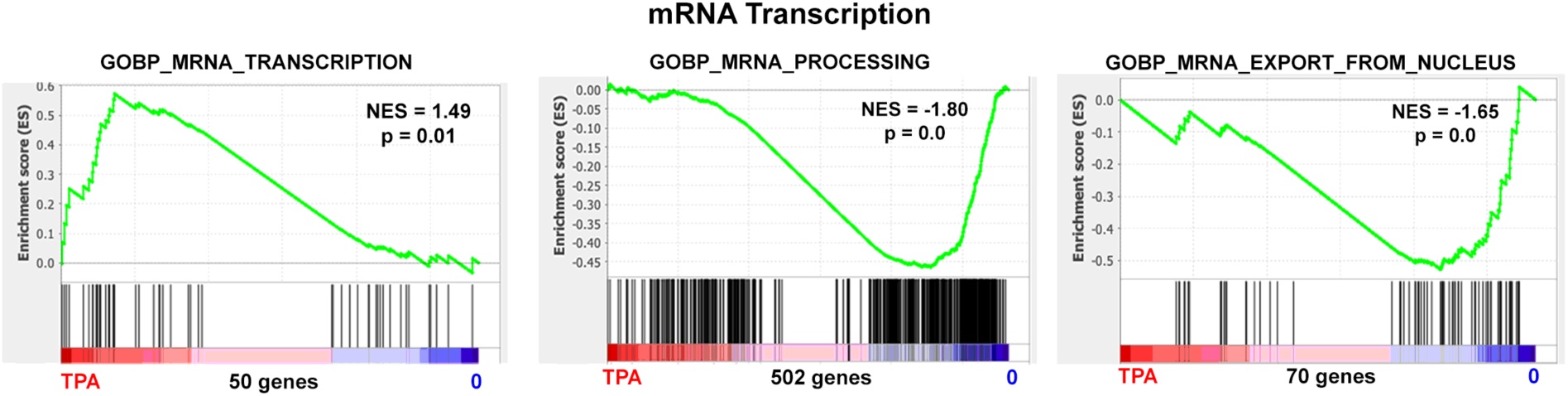
MRNA. GSEA plots tracking the progress of nascent mRNA from synthesis, to post-transcriptional modifications, to transport out of the cell nucleus into the cytoplasm for TPA induced HL-60/S4 macrophages. The latter two physiological steps indicate significant reductions in the necessary gene transcripts.

The final status of the “finished” mRNAs with polyA tails are presented in Table 2. “Differential Gene Expression of RA and TPA treated HL-60/S4 Cells” documents that not all genes exhibit increased relative (e.g., RA/0 and TPA/0) transcription. Some genes show increased DGE. Some are decreased; some are not changed. Table 2 was adapted from (Mark Welch et al., 2017) after modifications for splicing and mapping to hg38.

**Table 2.** Number of genes with significant changes in relative transcript levels for HL-60/S4 differentiation states. Column titles: DGE, Differential Gene Expression; RA, 4 days of retinoic acid differentiation to granulocyte form; TPA, 4 days of phorbol ester (TPA) differentiation to macrophage form. Total and Control are the number of genes mapped with normalized mean read count of at least 1. % is the percentage of the appropriate Total for the specific cell condition. For a complete list of all analyzed genes, see Table S1.

| Differential Gene Expression of RA and TPA treated<br>HL-60/S4 Cells |  |  |  |  |
| --- | --- | --- | --- | --- |
| DGE | RA count | RA % | TPA count | TPA % |
| Increased | 4107 | 23% | 4839 | 27% |
| Decreased | 3645 | 21% | 4236 | 24% |
| Unchanged | 9815 | 56% | 8785 | 49% |
| Total | 17567 |  | 17860 |  |
| Control | 17201 |  | 17203 |  |

### mRNA Translation: Downregulation in HL-60/S4 Macrophages

Apparently, the translational apparatus of HL-60 macrophages is less than optimal for all mRNAs. The HL-60/S4 macrophages exhibit dramatic reductions of transcripts for rRNAs and ribosomal proteins that are necessary for proper nucleolar function. This is documented in the following three GSEA plots of Figure 11: “GOCC_NUCLEOLUS”; “GOBP_RIBOSOME_BIOGENESIS”; “GOBP_RIBOSOME_ASSEMBLY”. The fact that SASP transcripts can succeed in being translated, while many other transcripts cannot, has been recognized before (Payea et al., 2021). Unfortunately, this paradox remains to be adequately explained. It may be that the major functions of macrophages (e.g., clearing tissues of infectious agents, cancer cells, damaged cells and facilitating wound healing) have been secured by evolutionary selection and depend upon the SASP protein products.

**Figure 11.**
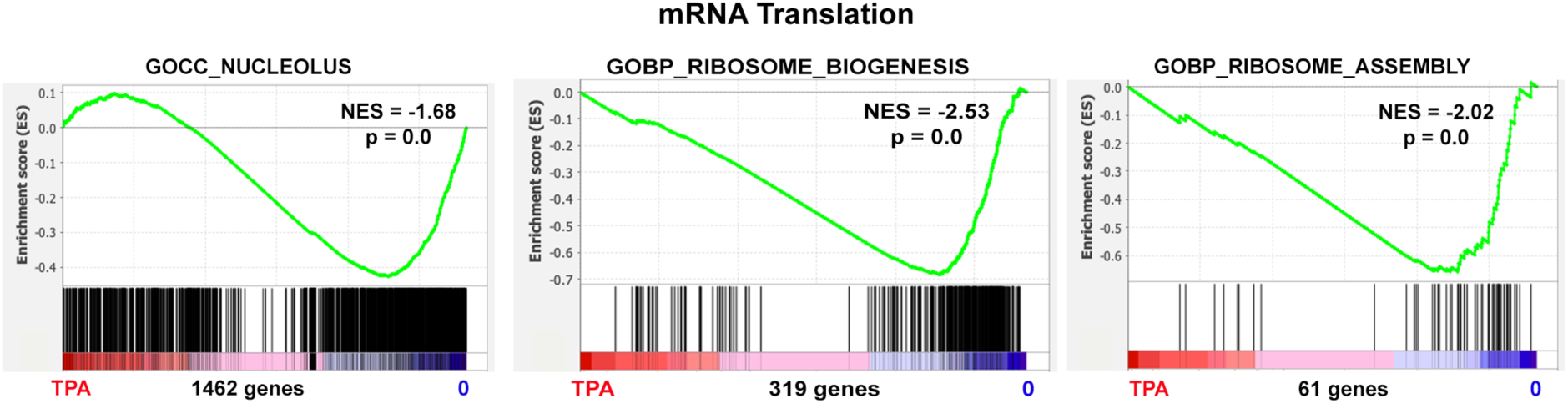
Translation. GSEA plots documenting the reduction of transcripts in HL-60/S4 macrophage related to nucleolus organization and ribosome formation, needed for translation of mature polyA-mRNA.

### HL-60/S4 Macrophages Exhibit Enriched Apoptosis Transcripts

There is a prevalent view that senescent cells resist apoptosis (Childs et al., 2014; Hu et al., 2022; Salminen et al., 2011). In an attempt to resolve this issue in the HL-60/S4 cell system, we performed GSEA employing two Apoptosis gene sets (Figure 12a). The analyses demonstrate that for each apoptosis gene set, there is significant enrichment of apoptosis genes in TPA treated cells. Figure 12b indicates the Log_2_FC (TPA/0) for these common apoptosis genes. The two gene sets display considerable overlap in genes comprising their Leading Edges. It is interesting that the three common genes with the greatest Log_2_FC > 0 are related to the TNF (Tumor Necrosis Factor) superfamily. We suggest that the HL-60/S4 cell system is a clear example of senescent cells that do <u>not</u> resist apoptosis. However, lack of resistance does not prove that HL-60/S4 macrophages die by apoptosis. This question is discussed later with data in *Inflammation/Inflammasomes in HL-60/S4 macrophage*

**Figure 12.**
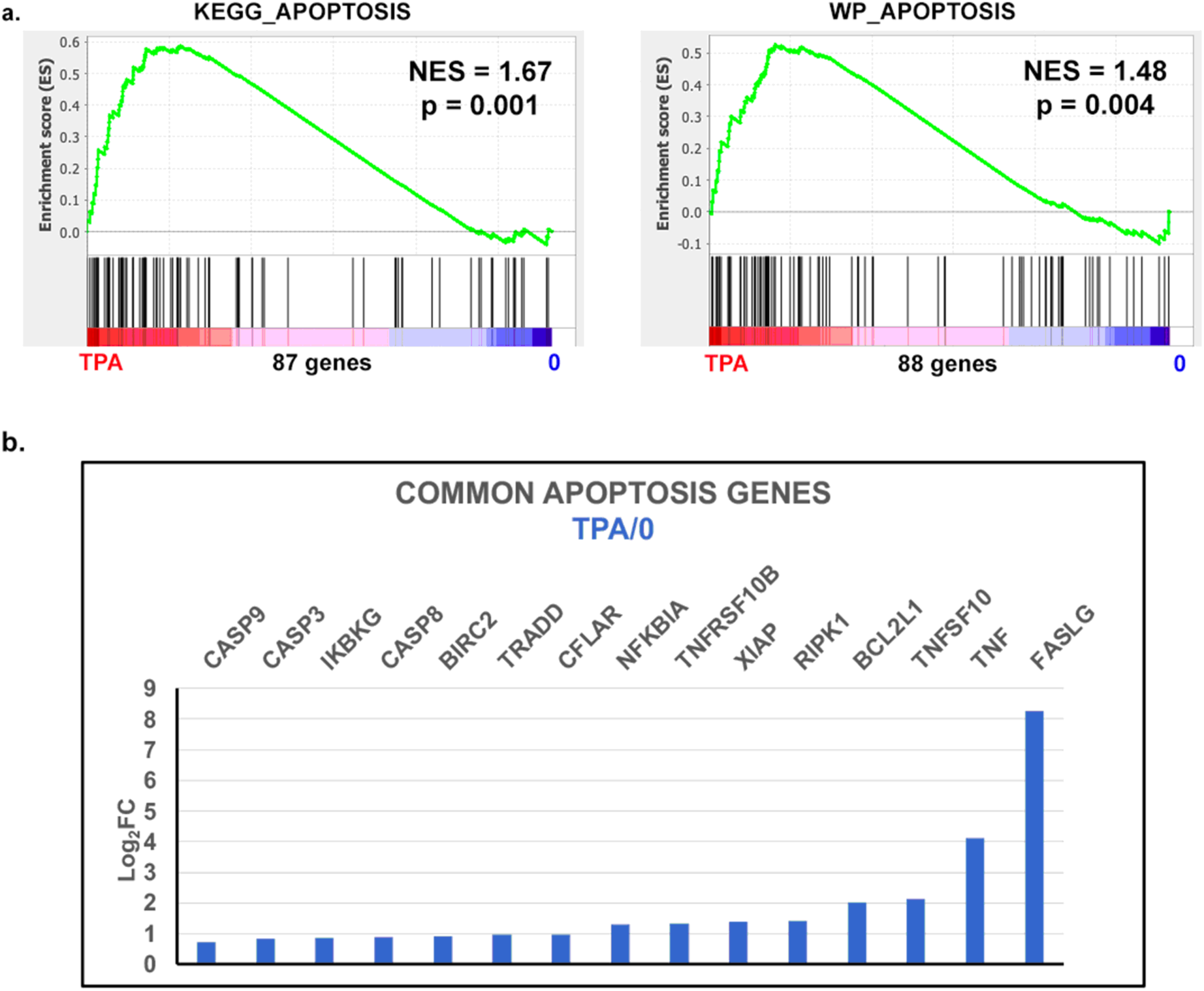
Apoptosis. a) GSEA plots for apoptosis enrichment in the TPA treated HL-60/S4 cells. The NES values for both gene sets are > 0; nominal p values are clearly significant (i.e., ∼0.00). b) Log_2_FC differential gene expression values for the common apoptosis genes in the two Leading Edges. It should be pointed out that RIPK1 participates in both apoptosis and necroptosis. In addition, BCL2L1 exists in two isoforms, one pro- and one anti-apoptosis. The solid bars indicate a PPDE > 0.95.

### Signaling Senescence: TGFB1 and NOTCH1

The previous references on the multiple origins and pathways to cellular senescence dramatically illustrate its complexity. In the HL-60/S4 system, the initial stress on the undifferentiated cells is treatment with TPA (phorbol ester). It has been known for many years that TPA mimics diacylglycerol (DAG), which is a “natural” activator of protein kinase C (PKC) in HL-60 cells (Ebeling et al., 1985; Yamamoto et al., 2006). More recently, it has been shown that PKCs (esp., PKC-δ or PRKCD) are downstream mediators of TGFB1-induced cellular senescence (Katakura et al., 2009). This applies to both replicative and oncogene-induced senescence. Among the many effects of TGFB1 towards senescence induction are the increase in transcription of the CDK inhibitors p16 and p21 (both observed, Table 1). In addition, there is evidence that TGFB1 enhances ROS (Reactive Oxygen Species) production, which can also be considered a stress (Tominaga and Suzuki, 2019). Furthermore, TGFB1 has the property of “autoinduction”; it can induce itself, leading to more prolonged and amplified effects on cells (Hariyanto et al., 2021).

The possible role of NOTCH1 in the induction of senescence has been explored in primary human endothelial cells (Venkatesh et al., 2011). During *in vitro* propagation of these primary cells, there is increased expression of NOTCH receptors and ligands. Senescence is induced at low passage, with increased expression of p21.

Within the past decade, there has been new thinking about NOTCH induced senescence (NIS) (Hoare et al., 2016; Hoare and Narita, 2017; Hoare and Narita, 2018; Teo et al., 2019). It was previously thought that senescence arises from specific stresses which yield non-cycling cells that secrete a defined set of cytokines and chemokines (SASP) under the control of NOTCH1. The new view is more dynamic with changing cell states and changing SASPs. This has been explored in the context of Ras-induced senescence (RIS), where in the initial state, the cells exhibit high levels of NOTCH1 and TGFB1 and low levels of CEBPB (C/EBPβ, a transcription factor for SASP), resulting in low levels of proinflammatory factors (e.g., IL1B and IL6). Subsequently, the senescent cells develop low levels of NOTCH1 and TGFB1 with high levels of CEBPB, resulting in high levels of proinflammatory factors (e.g., IL1B and IL6) (Hoare and Narita, 2017; Hoare and Narita, 2018). The most recent model (Teo et al., 2019) refers to primary and secondary senescence states, with cooperative changes between NOTCH and SASP levels.

Interestingly, the Ras-induced senescence (RIS) gene transcript changes are <u>not</u> what we see with TPA treated HL-60/S4 cells. Figure 13a and b (left panel) display one HL-60/S4 macrophage GSEA plot for TGFβ signaling and two plots for NOTCH signaling, indicating that the senescent TPA cells are enriched with the relevant gene set transcripts. Figure 13b (right panel) displays Log_2_FC differential expression (TPA/0) values for genes described earlier in RIS. Note that in HL-60/S4 macrophages TGFB1, NOTCH1, CEBPB, IL1A, IL1B and IL6 are all simultaneously upregulated. Also upregulated is PDGFA (a gene for a growth factor, <u>not</u> upregulated in RIS cells). Another difference between the RIS cells and TPA macrophage is the downregulation of JAG1 and HMGA1 (in a NOTCH1 complex) for RIS juxtacrine regulation (Parry et al., 2018). Whereas, in Figure 13b (right panel), TPA macrophages exhibit upregulation in JAG2, HEY1 and DELTA1 (DLK1). This comparison between RIS and TPA-macrophage transcriptomes may not be justified, however, since the “stress” of RIS is oncogenic; but the “stress” on HL-60/S4 cells is presently unknown (see Discussion).

**Figure 13.**
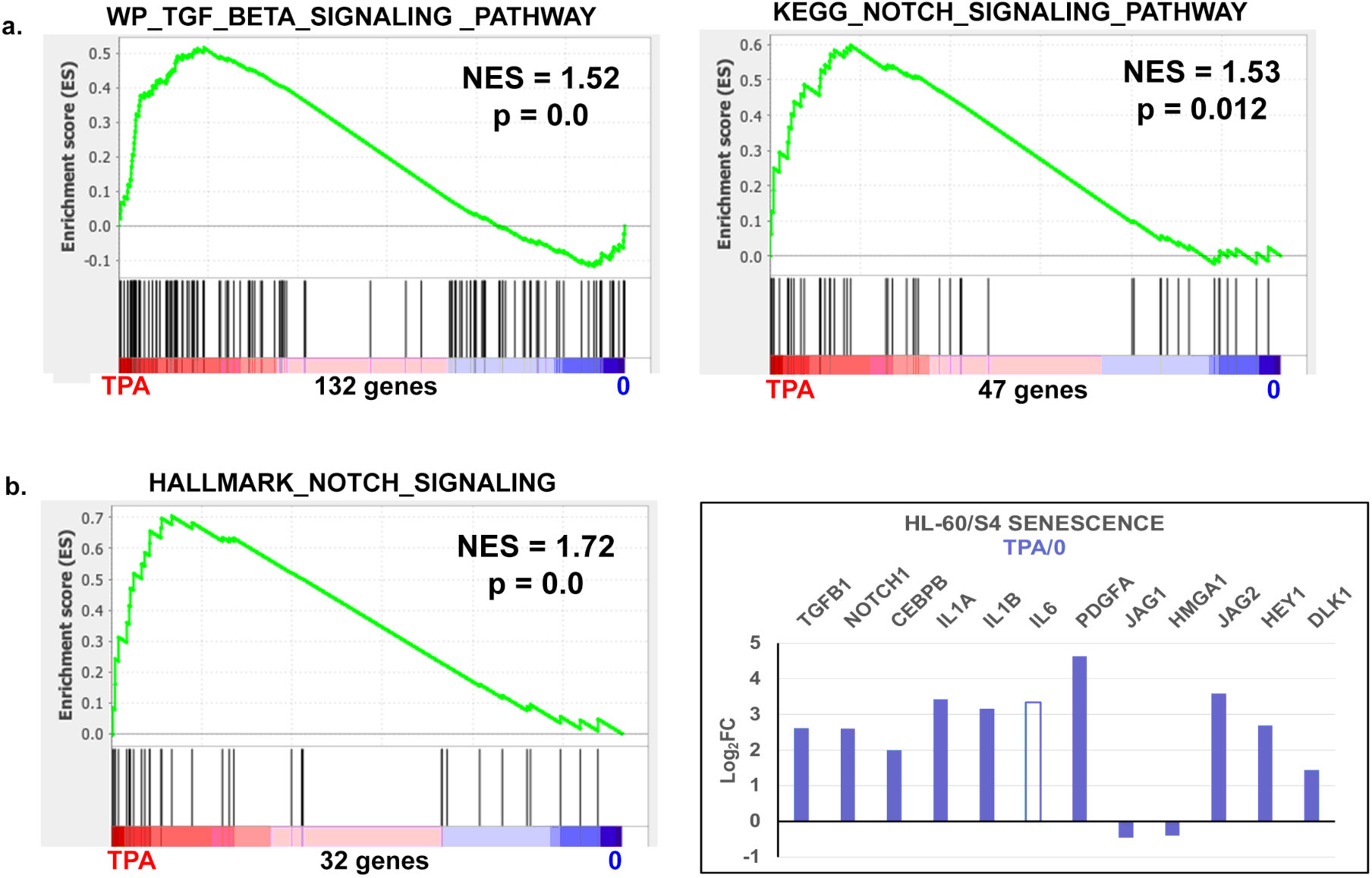
Signaling Senescence. a) and b) (left panel) GSEA plot for TGFβ signaling enrichment in the TPA treated HL-60/S4 cells and GSEA plots for NOTCH signaling enrichment in the TPA treated HL-60/S4 cells. The NES values for all three GSEA plots are significantly > 0.00. b) (right panel) Log_2_FC differential expression (TPA/0) values for the genes described earlier in RIS, but now examined in HL-60/S4 macrophages, including genes from the TGFB1 and NOTCH1 pathways. The open bar indicates lower statistical significance (PPDE < 0.95); solid bars indicate a PPDE > 0.95.

Even so, evidence favoring that HL-60/S4 macrophage are experiencing NOTCH induced senescence can be seen by examination of the two Leading Edges of the GSEA NOTCH plots: HALLMARK_NOTCH_SIGNALING and of KEGG_NOTCH_SIGNALING_PATHWAY presented in Table S2. This table demonstrates an increased expression of NOTCH receptors, ligands and processing enzymes.

### Oxidative Stress and DNA damage response in HL-60/S4 macrophages

Macrophages have a dangerous lifestyle. When pathogens or cell dangers are in the environment, macrophages synthesize and release ROS and RNS (i.e., reactive oxygen and reactive nitrogen species). These chemicals can also adversely affect macrophage cell metabolism, DNA integrity and viability. At the same time, macrophages can activate the anti-oxidant synthesis NRF2 pathway (Virág et al., 2019). The hydroxyl radical (.OH) is considered the most reactive ROS, able to damage proteins, lipids and DNA. Figure 12a presents one GSEA plot focused upon Oxidative Stress Response in HL-60/S4 macrophages. This plot clearly indicates that the respective gene sets are significantly enriched at the TPA end of the ranked gene distributions. The Leading Edge of WP_Oxidative_Stress_Response (Table S3) includes the gene NRF2 (alternate name in Table S1: NFE2L2), suggesting that the TPA macrophage has probably also activated the main pathway to alleviate oxidative stress. NRF2 is regarded as a major regulator of an anti-oxidation response and is also involved in control of autophagy (Ma, 2013). The Leading Edge also documents the increased transcription of anti-oxidation glutathione peroxidases and superoxide dismutase. In addition, Figure 14a presents two GSEA plots focused upon DNA Repair and DNA Damage Response. Both plots demonstrate that the DNA Repair and DNA Damage Response gene sets are <u>not</u> enriched at the TPA end of the ranked gene distributions. This implies that DNA Damage Repair (DDR) is not enhanced in the TPA induced macrophage, possibly because the NRF2 pathway is working properly. For comparison, Figure 14b presents a Log_2_FC graph of key DDR gene transcripts, identified in an important early paper on this subject (Broustas and Lieberman, 2014). Clearly, the DDR is not activated in HL-60/S4 macrophages.

**Figure 14.**
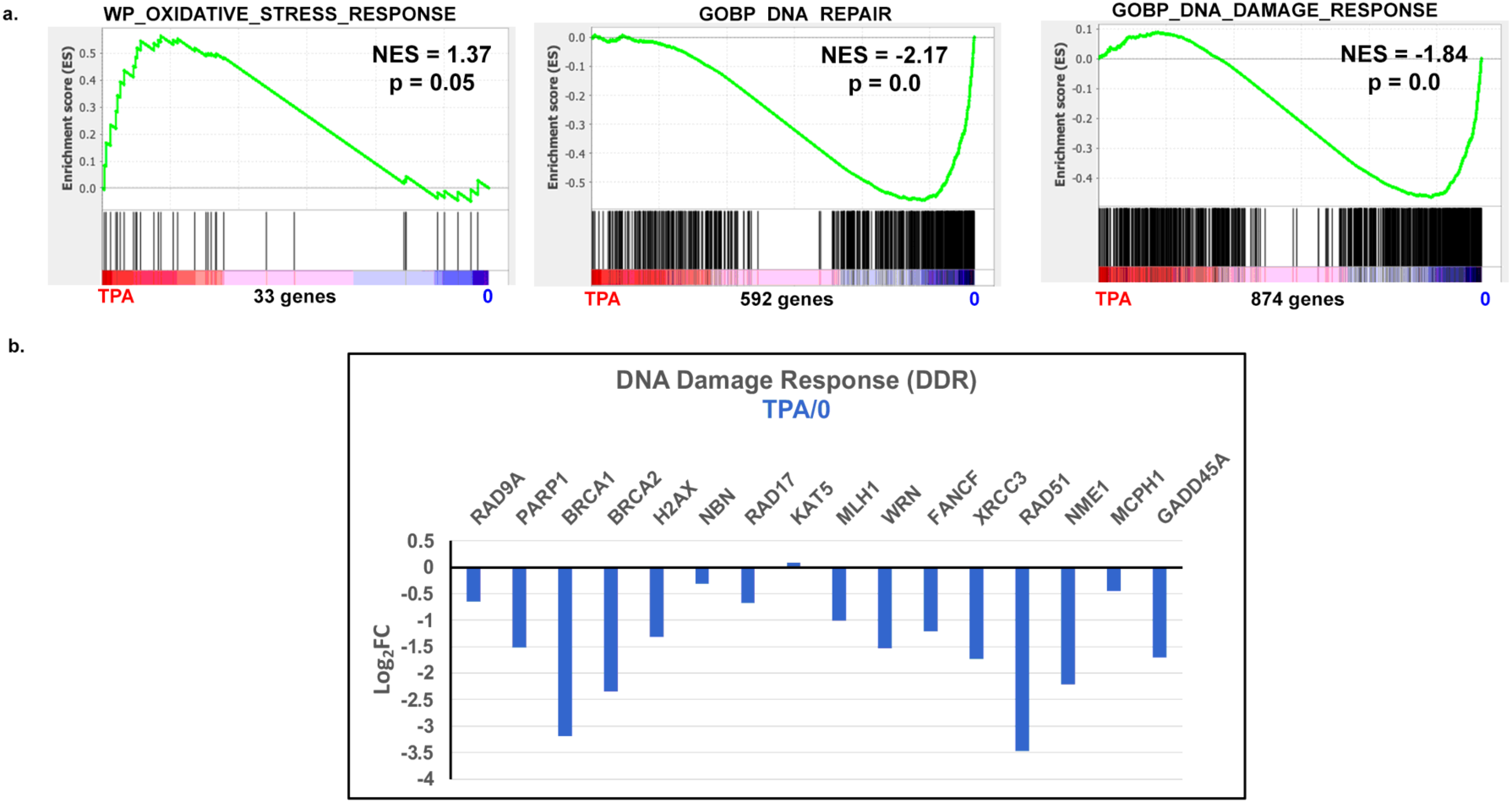
Oxidative Stress and DNA Damage. a) GSEA plots for Oxidative Stress Response, DNA Repair and DNA Damage Response in the HL-60/S4 macrophage. b) Log_2_FC differential expression (TPA/0) values for the DDR genes described earlier (Broustas and Lieberman, 2014), but now examined in the HL-60/S4 macrophage transcriptome. The solid bars indicate a PPDE > 0.95.

### Upregulation of Autophagy in the HL-60/S4 Macrophage

“Cellular Autophagy” is a catabolic process, involving degradation and recycling of cellular organelle components and macromolecules. Damaged organelles are enclosed within the membranes of an autophagosome, which is fused with a lysosome and digested by lysosomal enzymes. Autophagy has been identified in association with oncogenic-induced senescence (Narita et al., 2009). The coupling of autophagy and senescence has been recently reviewed (Cayo et al., 2021). Annotated gene lists for various stages of the autophagic pathway have been created, including lists amenable to GSEA (Bordi et al., 2021). A recent publication clearly demonstrates that HL-60 (i.e., the parent cell line for HL-60/S4) macrophage undergo autophagy (Mandic et al., 2022). Independently, we have arrived at a similar conclusion, employing GSEA (including a gene list from Bordi et al., 2021) on the transcriptome data from TPA treated HL-60/S4 cells. These analyses (Figures 15a and b [left side], employing gene sets from the GSEA website) substantiate that autophagy is present in the HL-60/S4 senescent macrophage. Figure 15b [right side] uses a gene set from the Autophagy “Induction List” (Bordi et al., 2021). Figure 15c displays Log_2_FC differential expression (TPA/0) values for the Autophagy “Induction List” genes examined in the HL-60/S4 macrophage (TPA/0) transcriptome. BCL2 is significantly downregulated more than 2-fold. Because BCL2 is considered to be anti-apoptotic, its downregulation agrees with our analysis (above) of apoptotic GSEA gene sets. We propose that apoptosis and autophagy can both occur in the HL-60/S4 senescent macrophage. The question of competition between apoptosis and autophagy has been considered in an excellent review (Fan and Zong, 2013). From an evolutionary point-of-view, it is interesting that autophagy is considerably “older” (e.g., being present in unicellular yeast); apoptosis appeared later, during the evolution of metazoans.

**Figure 15.**
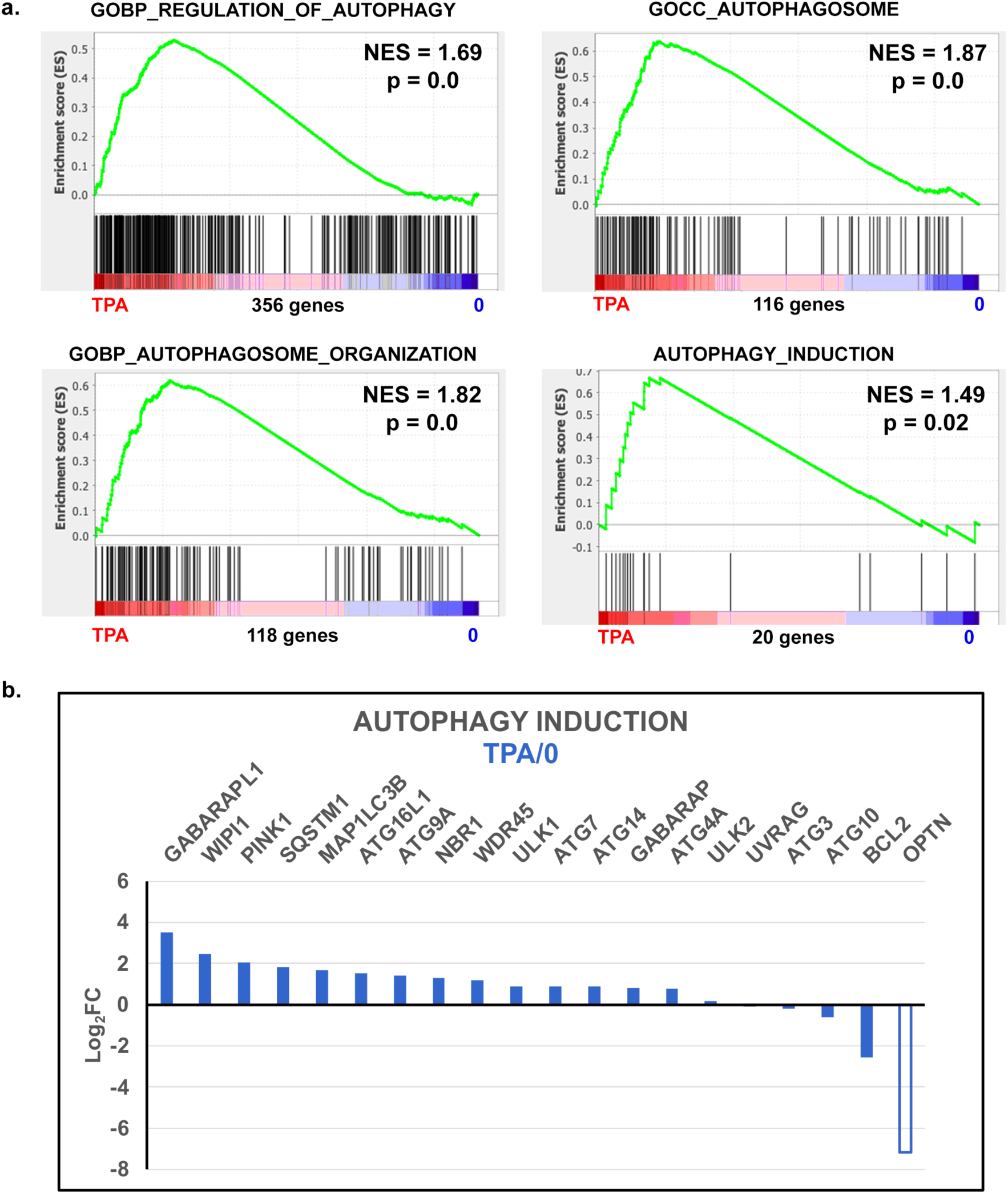
Autophagy. a) GSEA plots for Regulation of Autophagy, Autophagosome, Autophagosome Organization and for Autophagy Induction in the HL-60/S4 macrophage. b) Log_2_FC differential expression (TPA/0) values for Autophagy Induction in the HL-60/S4 macrophage. The open bar indicates lower statistical significance (PPDE < 0.95); solid bars indicate a PPDE > 0.95.

### The Extracellular Matrix (ECM) of the HL-60/S4 Macrophage

The ECM is a network of polymers of varying composition that depend upon the cell or tissue-type (Yue, 2014). The composition and structure are dynamic. It may attach the cells to a substrate and/or other cells, and it may participate in storing bioactive molecules. The protein building blocks of an ECM may include collagen, proteoglycans, laminin, fibronectin and elastin, among others. We present GSEA results for two MSigDB gene sets matched against the transcriptomes of TPA treated HL-60/S4 cells in Figure 16a. Log_2_FC differential expression values for selected genes from the Leading Edge of both plots are presented in Figure 16b. There is a clear enrichment of bioactive TGFβ, NOTCH1 and ECM building blocks in TPA treated HL-60/S4 cells.

**Figure 16.**
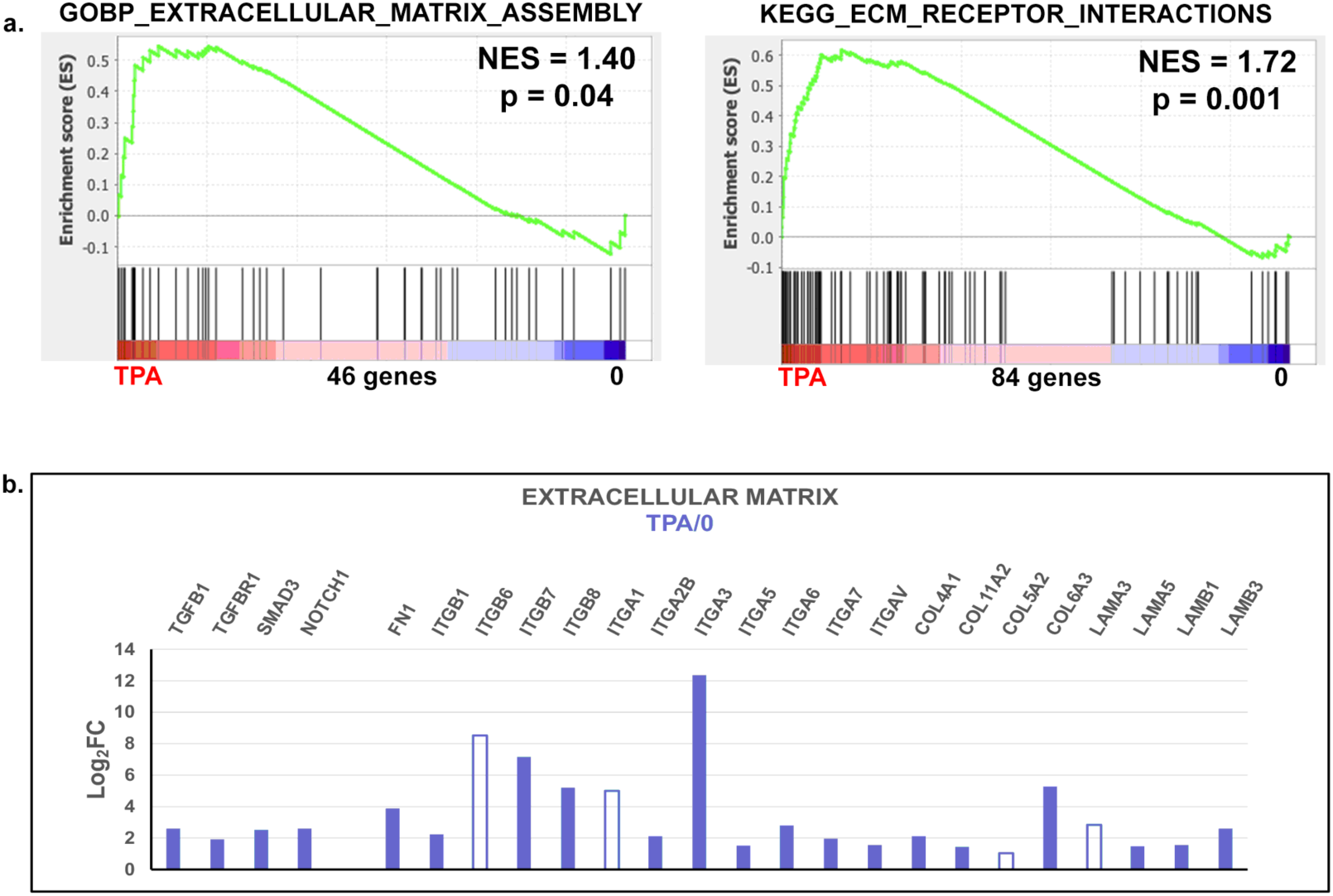
a) GSEA plots for Extracellular Matrix Assembly (left side) and ECM Receptor Interaction (right side) in the TPA treated HL-60/S4 macrophage. b) Log_2_FC differential expression (TPA/0) values for selected genes from the Leading Edges in both plots of the HL-60/S4 macrophage (Part a). The left four bars are from the left plot of (Part a); the right twenty bars are from the right plot of (Part a). The open bars indicate lower statistical significance (PPDE < 0.95); solid bars indicate a PPDE > 0.95.

### Inflammation/Inflammasomes in HL-60/S4 macrophage

One of the critical phenotypic questions is whether the HL-60/S4 macrophage can sustain an inflammatory response. Since this study is focused upon *in vitro* cell culture, rather than *in vivo* animal tissues, the criteria must be defined at the cellular level (e.g., secretion of inflammatory chemokines and cytokines). Another criterion is the production of activated inflammasomes (Guo et al., 2015; Swanson et al., 2019; Ta and Vanaja, 2021; Zheng et al., 2020). Figure 17a presents GSEA plots of appropriate gene sets to establish that HL-60/S4 macrophages exhibit inflammatory responses during the differentiation period. Figure 17b (left side) presents a GSEA plot showing that HL-60/S4 macrophages produce inflammasomes. Figure 17b (right side) shows a GSEA plot demonstrating that HL-60/S4 macrophages succumb to Pyroptosis, a form of caspase-mediated inflammatory cell death (Robinson et al., 2019). Figure 17c presents Log_2_FC differential expression (TPA/0) values for the Leading Edge of the Pyroptosis GSEA plot. It is important to point out that the current perspective is that Pyroptosis is an inflammatory process of programmed cell death; whereas, apoptosis is regarded as a non-inflammatory process of programmed cell death (Ketelut-Carneiro and Fitzgerald, 2022).

**Figure 17.**
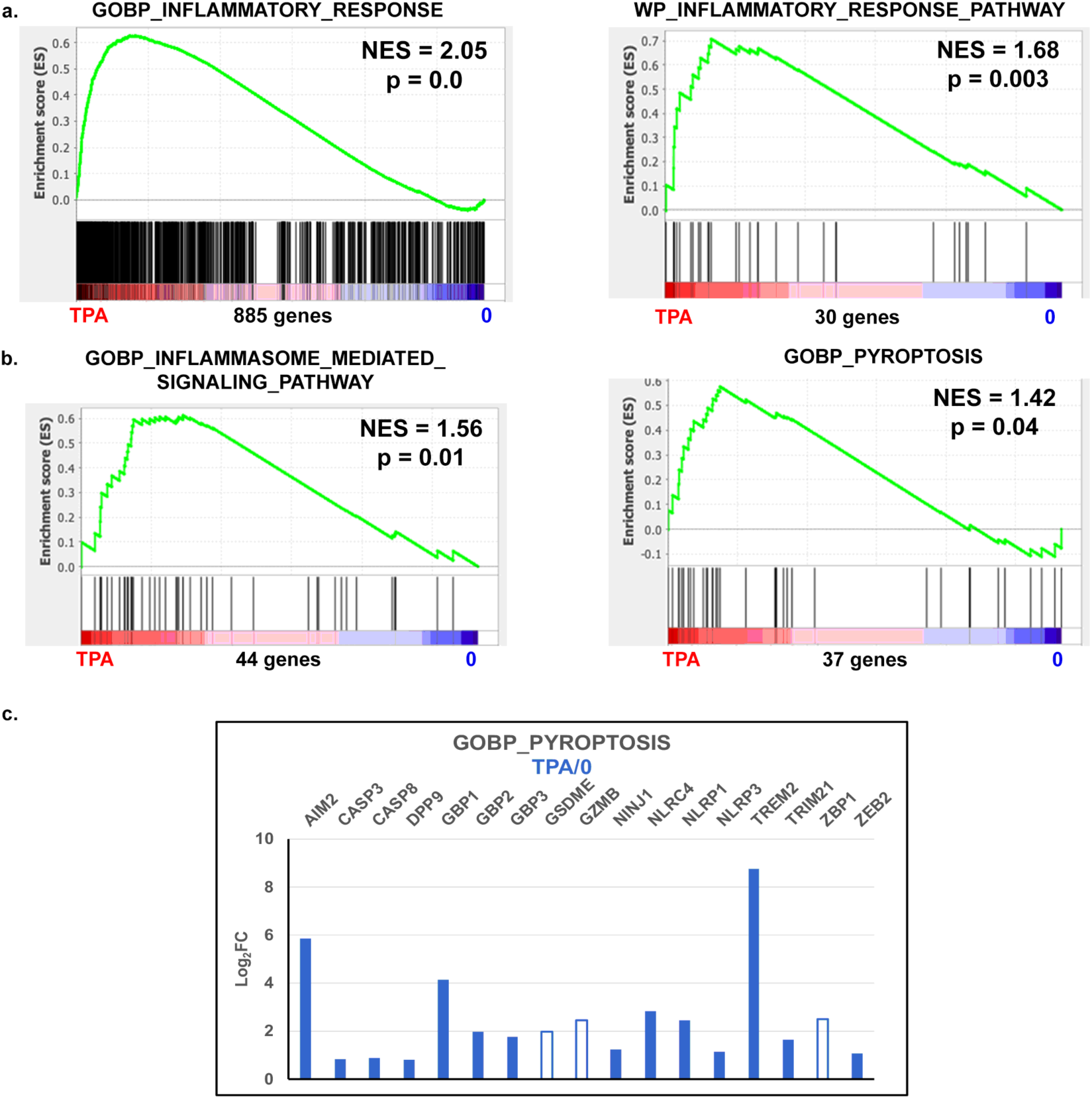
a) GSEA plots for Inflammatory Response and Inflammatory Response Pathway. b) GSEA plots for Inflammasome Mediated Signaling Pathway and for Pyroptosis in TPA treated HL-60/S4 macrophages. c) Log_2_FC differential expression (TPA/0) values for Pyroptosis genes in the Leading Edge. The open bars indicate lower statistical significance (PPDE < 0.95); solid bars indicate a PPDE > 0.95.

An additional level of complexity in interpreting the data and conclusions of this publication comes from asking the question: What is the “polarity” (class) of TPA differentiated HL-60/S4 macrophages? A simplistic form of this question becomes: M1 macrophages are pro-inflammatory; M2 macrophages are anti-inflammatory. Is the HL-60/S4 TPA macrophage M1 or M2? Our oversimplified answer is: They resemble M1 somewhat more than M2. This conclusion is based primarily upon searching the HL-60/S4 macrophage transcriptome for secretions and cell surface markers, that are described as being characteristic for the two classes of cells (Apeku et al., 2024; Strizova et al., 2023; Viola et al., 2019). In addition, a recent study has concluded that M1 macrophages can die by Pyroptosis (Li et al., 2022). Nonetheless, a more definitive study on the “M status” of the HL-60/S4 macrophage is desirable. Such a study must contend with the increased complexity of M2 subclasses and the acknowledged “plasticity” of macrophage polarity (Sica and Mantovani, 2012; Smith et al., 2016).

## Discussion

The purpose of this study is to characterize the various facets that constitute the multifaceted phenotype of a particular cell state; i.e., the TPA differentiated senescent macrophages of the HL-60/S4 cell line. We chose this cell state because: 1) It is clearly senescent. 2) Senescence is reproducibly and easily induced by the addition of TPA for 4 days. 3) It is recognizably different than other cell states of HL-60/S4, when viewed in the microscope. 4) We have detailed expression data for each of the HL-60/S4 cell states (i.e., 0, RA and TPA), offering a unique molecular view of the senescent cell.

The backbone of this study has been to employ GSEA (Gene Set Enrichment Analysis) to describe particular phenotypic facets, generating a “normalized enrichment analysis” (NES) with low nominal p values. This type of study is limited by the number and quality of available gene sets. That is the reason why multiple related gene sets were frequently employed in this study and compared in their Leading Edge, searching for significant similarities and differences. Specific gene enrichment at the TPA end of the “ranked genes” is equivalent to saying that the specific phenotypic facet is “upregulated”. Table 3 presents a summary classification of the various analyzed phenotypic facets into two categories: Gene set enrichment (NES > 0); Gene set reduction (NES < 0).

**Table 3.** Phenotypic Facets of HL-60/S4 Macrophages. Column Headings: GENE SET ENRICHMENT (NES>0), upregulation or increased gene transcription; GENE SET REDUCTION (NES<0), downregulation or decreased gene transcription.

| GENE SET ENRICHMENT<br>(NES>0) | GENE SET REDUCTION<br>(NES<0) |
| --- | --- |
| Senescence | -- |
| -- | Heterochromatin Changes |
| -- | Nucleosome Formation |
| -- | Chromatin Remodeling |
| mRNA Transcription | -- |
| -- | mRNA Translation |
| Apoptosis | -- |
| TGFB1 and NOTCH1 Signaling | -- |
| Oxidative Stress | DNA Damage and Repair |
| Autophagy | -- |
| Extracellular Matrix | -- |
| Inflammation/Inflammasomes | -- |

The utility of Table 3 is that it suggests broad research questions concerning possible relationships between HL-60/S4 macrophage phenotypic characteristics. For example: Why does the increased transcription of genes related to “Oxidative Stress and Damage” <u>not</u> lead to increased transcription of genes related to “DNA damage and repair”? We speculate that the oxidative damage was to cellular proteins and/or lipids, and not to the nuclear DNA. Another example: What is the basis of the “loss” of heterochromatin within the senescent HL-60/S4 macrophage nucleus? We speculate that a reduced transcription of some histones (e.g., H1 and H2A) and HP1 proteins leads to a reduction of chromatin higher-order structure. Changes in epigenetic histone modifications may also contribute.

In addition to employing GSEA to describe the phenotype of HL-60/S4 macrophage, we submitted the TPA/0 transcriptome to WebGestalt (https://www.webgestalt.org/option.php), for over-representation of KEGG pathways. We submitted 4714 genes with significantly increased transcript levels (PPDE > 0.95). The most over-represented KEGG pathway was (surprisingly) “Osteoclast Differentiation”. Exploring the KEGG pathway gene set list, we discovered the cell membrane TNF superfamily receptor TNFRSF11A, otherwise known as RANK, which is activated by RANKL (Huang et al., 2017), a combination essential for osteoclast differentiation. This observation opens the possibility of *in vitro* differentiation of HL-60/S4 cells into osteoclasts.

Another interesting question that can be explored using the HL-60/S4 cell system is whether the acquisition of senescence depends upon the type of substrate (or ECM) coating the microscope slide (e.g., polylysine, fibronectin or slides scraped of prior cell growth). We want to see whether properties (e.g., cell and nuclear shape) of 0, RA and TPA cells are altered by the nature of the substrate.

A very important fundamental question is: What is the stress that initiates development of senescence in the HL-60/S4 macrophages? As mentioned earlier the addition of TPA to undifferentiated HL-60/S4 cells mimics the normal differentiation response of added diacylglycerol (DAG). We suggest a hypothesis that encompasses many of the phenotypic characteristics listed in Table 3, based largely on GSEA results, and partly based upon prior publications (Behmoaras and Gil, 2021; Cayo et al., 2021; Robinson et al., 2019).

The hypothesis:

I) The stress is <u>programmed</u> macrophage differentiation.
II) This differentiation path leads to cellular senescence, probably both TGFβ and Notch-induced, including a halted cell cycle at the G1/S boundary and a SASP with pro-inflammatory secretions.
III) Autophagy is upregulated as a mechanism to degrade (oxidative) damaged molecules and organelles, and as an intracellular molecular recycling mechanism.
IV) Senescent macrophages appear to be dormant, i.e., inactive, waiting for induction of an inflammatory response. There may be cellular heterogeneity during dormancy, e.g., heterogeneity of pro- and anti-inflammatory state and composition of the SASP.
V) After the inflammatory response, senescent macrophages likely die by programmed cell death (e.g., apoptosis and/or pyroptosis) involving caspases.

A fundamental question pertaining to the present study is: How are the numerous phenotypic facets (e.g., the “conclusions” from individual gene set enrichment analyses) of the HL-60/S4 macrophages integrated within an individual cell. Are these phenotypic facets interdependent on one-another, or are they independent? Intracellular connectivity or independence would have been selected by evolution. Is this multiplicity of phenotypic facets observed in primary macrophages? There is a relevant parallel to the above questions, described in a recent review of Apoptosis, Pyroptosis and Necroptosis pathways (Ketelut-Carneiro and Fitzgerald, 2022). The authors write: “Although they are distinct cell death pathways, apoptosis, pyroptosis, and necroptosis constitute a single, synchronized and coordinated cell death system in which any one pathway can compensate for another, or all three can work in the same cell, but at different moments depending on the context and the time.” We suggest that a similarly evolved interdependence connects numerous phenotypic facets within a single HL-60/S4 senescent macrophage cell.

## Acknowledgements

ALO and DEO express our appreciation to the University of New England for allowing us to use laboratory space for the past 11 years. We also express our gratitude to the MaineHealth Institute for Research for allowing us to join the research group of Dr. Igor Prudovsky.

