## Supplemental Table 2 for "The Multifaceted Phenotype of Senescent HL-60/S4 Macrophages"

**Table S2**

**LEADING EDGES OF GSEA (TPA/0) PLOTS**

| **HALLMARK_NOTCH_SIGNALING** | **KEGG_NOTCH_SIGNALING_PATHWAY** |
| --- | --- |
| [ARRB1](https://ensembl.org/Search/Results?q=ARRB1) | [ADAM17](https://ensembl.org/Search/Results?q=ADAM17) |
| [CCND1](https://ensembl.org/Search/Results?q=CCND1) | [CREBBP](https://ensembl.org/Search/Results?q=CREBBP) |
| [DTX1](https://ensembl.org/Search/Results?q=DTX1) | [DTX1](https://ensembl.org/Search/Results?q=DTX1) |
| [DTX2](https://ensembl.org/Search/Results?q=DTX2) | [DTX2](https://ensembl.org/Search/Results?q=DTX2) |
| [FBXW11](https://ensembl.org/Search/Results?q=FBXW11) | [DTX3L](https://ensembl.org/Search/Results?q=DTX3L) |
| [FZD7](https://ensembl.org/Search/Results?q=FZD7) | [DVL3](https://ensembl.org/Search/Results?q=DVL3) |
| [LFNG](https://ensembl.org/Search/Results?q=LFNG) | [EP300](https://ensembl.org/Search/Results?q=EP300) |
| [MAML2](https://ensembl.org/Search/Results?q=MAML2) | [JAG2](https://ensembl.org/Search/Results?q=JAG2) |
| [NOTCH1](https://ensembl.org/Search/Results?q=NOTCH1) | [KAT2B](https://ensembl.org/Search/Results?q=KAT2B) |
| [NOTCH3](https://ensembl.org/Search/Results?q=NOTCH3) | [LFNG](https://ensembl.org/Search/Results?q=LFNG) |
| [PPARD](https://ensembl.org/Search/Results?q=PPARD) | [MAML1](https://ensembl.org/Search/Results?q=MAML1) |
| [PRKCA](https://ensembl.org/Search/Results?q=PRKCA) | [MAML2](https://ensembl.org/Search/Results?q=MAML2) |
| [PSEN2](https://ensembl.org/Search/Results?q=PSEN2) | [MAML3](https://ensembl.org/Search/Results?q=MAML3) |
| [ST3GAL6](https://ensembl.org/Search/Results?q=ST3GAL6) | [NCSTN](https://ensembl.org/Search/Results?q=NCSTN) |
| [TCF7L2](https://ensembl.org/Search/Results?q=TCF7L2) | [NOTCH1](https://ensembl.org/Search/Results?q=NOTCH1) |
|  | [NOTCH2](https://ensembl.org/Search/Results?q=NOTCH2) |
|  | [NOTCH3](https://ensembl.org/Search/Results?q=NOTCH3) |
|  | [NUMB](https://ensembl.org/Search/Results?q=NUMB) |
|  | [PSEN1](https://ensembl.org/Search/Results?q=PSEN1) |
|  | [PSEN2](https://ensembl.org/Search/Results?q=PSEN2) |
|  | [PTCRA](https://ensembl.org/Search/Results?q=PTCRA) |

Genes in each column are arranged in alphabetical order.
