## Supplemental Table 3 for "The Multifaceted Phenotype of Senescent HL-60/S4 Macrophages"

**Table S3**

**LEADING EDGE OF GSEA (TPA/0) PLOT**

**WP_OXIDATIVE_STRESS_RESPONSE**

[CYBB](https://ensembl.org/Search/Results?q=CYBB)

[FOS](https://ensembl.org/Search/Results?q=FOS)

[GCLC](https://ensembl.org/Search/Results?q=GCLC)

[GPX1](https://ensembl.org/Search/Results?q=GPX1)

[GPX3](https://ensembl.org/Search/Results?q=GPX3)

[GSR](https://ensembl.org/Search/Results?q=GSR)

[HMOX1](https://ensembl.org/Search/Results?q=HMOX1)

[JUNB](https://ensembl.org/Search/Results?q=JUNB)

[MAPK14](https://ensembl.org/Search/Results?q=MAPK14)

[NFE2L2](https://ensembl.org/Search/Results?q=NFE2L2)

[NOX5](https://ensembl.org/Search/Results?q=NOX5)

[NQO1](https://ensembl.org/Search/Results?q=NQO1)

[SOD3](https://ensembl.org/Search/Results?q=SOD3)

[UGT1A6](https://ensembl.org/Search/Results?q=UGT1A6)

Genes are arranged in alphabetical order.
